# Transcriptome-wide characterization of genetic perturbations

**DOI:** 10.1101/2024.07.03.601903

**Authors:** Ajay Nadig, Joseph M. Replogle, Angela N. Pogson, Steven A McCarroll, Jonathan S. Weissman, Elise B. Robinson, Luke J. O’Connor

**Author notes:** Correspondence (A.N.), (L.J.O), (J.M.R.).

## Abstract

Single cell CRISPR screens such as Perturb-seq enable transcriptomic profiling of genetic perturbations at scale. However, the data produced by these screens are often noisy due to cost and technical constraints, limiting power to detect true effects with conventional differential expression analyses. Here, we introduce TRanscriptome-wide Analysis of Differential Expression (TRADE), a statistical framework which estimates the transcriptome-wide distribution of true differential expression effects from noisy gene-level measurements. Within TRADE, we derive multiple novel, interpretable statistical metrics, including the “transcriptome-wide impact”, an estimator of the overall transcriptional effect of a perturbation which is stable across sampling depths. We analyze new and published large-scale Perturb-seq datasets to show that many true transcriptional effects are not statistically significant, but detectable in aggregate with TRADE. In a genome-scale Perturb-seq screen, we find that a typical gene perturbation affects an estimated 45 genes, whereas a typical essential gene perturbation affects over 500 genes. An advantage of our approach is its ability to compare the transcriptomic effects of genetic perturbations across contexts and dosages despite differences in power. We use this ability to identify perturbations with cell-type dependent effects and to find examples of perturbations where transcriptional responses are not only larger in magnitude, but also qualitatively different, as a function of dosage. Lastly, we expand our analysis to case/control comparison of gene expression for neuropsychiatric conditions, finding that transcriptomic effect correlations are greater than genetic correlations for these diagnoses. TRADE lays an analytic foundation for the systematic comparison of genetic perturbation atlases, as well as differential expression experiments more broadly.

## Introduction

A foundational approach in modern biology involves measuring the phenotypic effect of genetic variation, naturally occurring or experimentally induced, on cells. One of the most informative and scalable strategies for measuring cellular responses is *differential expression*, which quantifies RNA abundance for all genes by gene expression microarray or RNA sequencing. Recent technological advances have combined single-cell RNA sequencing with CRISPR screening to enable massively scalable transcriptomic profiling of genetic perturbations, in an approach called Perturb-Seq (Dixit et al. 2016; Adamson et al. 2016; Jaitin et al. 2016). Despite their promise, Perturb-seq screens produce data with variable amounts of estimation error across perturbations, and thus pose a challenge for conventional analytic methods, including differential expression and correlation, which generate variable results depending on statistical power. The extent to which this limitation has confounded understanding and comparison of perturbation experiments is unclear.

The field of human genetics has contended with a similar issue in genetic association studies - where power is typically limited due to finite sample sizes and small effect sizes - by estimating population parameters directly, without the use of significance thresholds. This approach is widely used to infer the total genetic effect (“SNP-heritability”; Yang et al. 2010), to identify disease-relevant cell types and pathways (“heritability enrichment”; Finucane et al. 2018), and to understand the shared genetic basis of different traits (“genetic correlation”; Bulik-Sullivan et al. 2015). A strength of this approach is that it distinguishes properties of a study or experiment from those of a trait or population.

In RNA-seq analysis, an analogous approach would be to estimate the *distribution of differential expression effects*, including those that are underpowered in a study, rather than testing for gene-wise differential expression. Some existing methods have attempted to go beyond significance thresholds to capture aspects of this distribution. The energy distance quantifies the strength of a perturbation as the difference between average between-condition vs. within-condition variability after normalization, filtering, and projection onto principal components (Replogle et al. 2022; Peidli et al. 2024). Gene-set enrichment analysis uses a rank-based approach to test gene-set enrichments of differential expression effects (Subramanian et al. 2005). Rank-rank hypergeometric overlap uses a similar approach to test for a significant correlation between differential expression effects across experiments *(Plaisier et al. 2010)*. *iDEA* uses a point-normal model for the distribution of differential expression effects to increase association power for the identification of individual differentially expressed genes (Ma et al. 2020). However, none of these approaches explicitly estimate and interpret the distribution of differential expression effects, and thus do not fully characterize of the transcriptome-wide consequences of perturbations.

Here, we present TRADE (TRanscriptome-wide Analysis of Differential Expression), a suite of statistical tools for formally modeling distributions of differential expression effects from RNA-seq experiments, including Perturb-seq. TRADE fits a flexible mixture model to estimated effects and standard errors to estimate the distribution of true differential expression effects. From this estimated distribution, we derive several highly interpretable metrics, including the transcriptome-wide impact, the effective number of differentially expressed genes, gene set enrichments, and correlation. We use TRADE to estimate and interpret these features for tens of thousands of genetic perturbations across two new, and three existing massive Perturb-Seq datasets (Replogle et al. 2022). Finally, we use TRADE to compare the effects of perturbations across cell types, to estimate dose-response curves for transcriptome-wide effects, and to estimate the bivariate transcriptomic relationships between neuropsychiatric conditions.

## Results

### Overview of methods

Consider an RNA-seq experiment comparing two conditions (e.g., perturbed and unperturbed). A conventional differential expression analysis fits a generalized linear model for each gene to estimate the difference in mean expression between conditions, producing a point estimate of the log_2_(Fold Change) and a standard error or p-value. The point estimate for gene *g* can be modeled as the sum of a true effect, β_*g*_, and a residual, ε_*g*_:

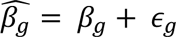

TRADE is a method to estimate the distribution of β from that of 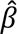 accounting for sampling variation. This approach distinguishes properties of a cellular response from those of an experiment, such as sample size and sequencing depth.

TRADE takes as input differential expression point estimates and standard errors, which we presently compute using *DESeq2* (Methods; Love, Huber, and Anders 2014) applied to pseudo-bulk RNA-seq read count matrices (**Figure 1A**). TRADE then estimates the distribution of β by fitting a mixture model to the distribution of effect size estimates, incorporating standard errors to account for sampling variation, using *ash (***Methods**; Stephens 2016). While *ash* was initially designed to perform Empirical Bayes shrinkage using the estimated effect size distribution as a prior, we instead focus on interpreting that distribution itself.

**Figure 1:**
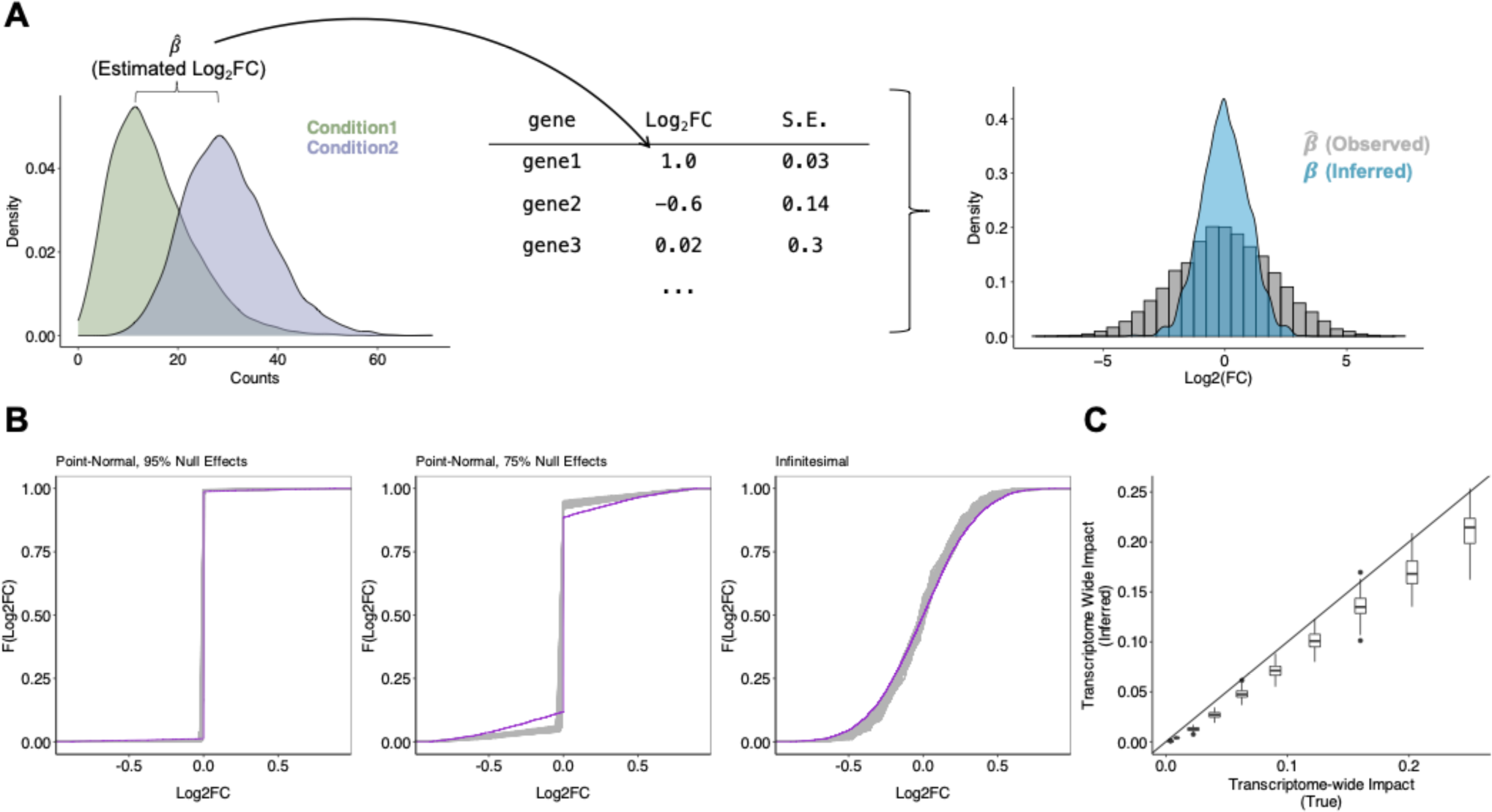
Transcriptome-wide Analysis of Differential Expression. **(A)** Schematic for TRADE analysis, starting from condition-wise gene expression counts and ending with estimated distribution of Log2FC. **(B)** Estimation of various simulated effect size distributions (Point-Normal with 95% Null, Point-Normal with 75% Null, Infinitesimal/normally distributed). Purple trace shows true effect size distribution; gray traces show estimated distributions across 100 replicates. **(C)** Comparison of estimated and true transcriptome-wide impact in simulations.

From this inferred effect distribution, TRADE computes several interpretable features that describe the transcriptome-wide landscape of differential effects. One key estimand is Var(β), the variance of the effect size distribution, which we term the “transcriptome-wide impact” (TI) (**Methods**). The transcriptome-wide impact can be interpreted as a measure of the overall transcriptomic change in a differential expression experiment, where perturbations with large transcriptome-wide impact include those with large effects on a few genes as well as those with smaller effects on many genes. TRADE also derives other estimates from the effect-size distribution, including gene-set enrichments, correlations between perturbations or cell types, and a novel measure of the effective number of differentially expressed genes, all of which are unbiased at finite sample size (**Methods**).

We tested our approach in simulations of varying effect size distributions, ranging from very sparse point-normal to fully infinitesimal, with a sample size of 200 cells per condition (**Methods; Figure 1B**). The inferred distributions generally differed from the true distributions only in the density around zero, by slightly overestimating the density at zero. Estimates of the transcriptome-wide impact (**Figure 1C**) were robust, with slight downward bias. The downward bias results from inadequate power to detect very small effects at finite sample size, even aggregating power across genes; at increased sample size, the bias disappears (**Supplementary Figure 1**).

### Transcriptome-wide impact of 9,866 genetic perturbations

We next sought to investigate the transcriptome-wide impact of a comprehensive set of genetic perturbations with TRADE. Recently, Repogle et al (2022) performed genome-scale Perturb-Seq with CRISPR interference (CRISPRi), which inhibits target gene transcription by recruitment of a dCas9-linked repressive KRAB domain, generating three datasets: K562-GenomeWide (perturbations of all 9,866 expressed genes in the K562 chronic myelogenous leukemia cell line), K562-Essential (2,057 common essential gene perturbations in the same cell line), and RPE1-Essential (2,393 common essential gene perturbations in a retinal pigmented epithelium cell line). To enable a more thorough comparison across cell types, we performed two additional large-scale Perturb-seq experiments targeting common essential genes in Jurkat and HepG2 cell lines: Jurkat-Essential (2,393 essential gene perturbations in a T-cell leukemia cell line) and HepG2-Essential (2,393 essential gene perturbations in a hepatocellular carcinoma cell line) (**Methods**). Key features of these datasets are summarized in **Table 1**.

**Table 1.** Characteristics of Perturb-Seq datasets analyzed.

| **Dataset**<br>**(Technology)** | **Reference** | **Cell**<br>**type** | **Perturbed**<br>**Gene-Set** | **Number of**<br>**perturbations** | **Median #**<br>**cells per**<br>**perturbation** |
| --- | --- | --- | --- | --- | --- |
| K562-GenomeWide<br>(CRISPRi) | Replogle et al,<br>2022, | K562 | All expressed<br>genes | 9866 genes | 178 |
| K562-Essential<br>(CRISPRi) | Replogle et al,<br>2022 | K562 | Essential genes | 2057 | 121 |
| RPE1-Essential<br>(CRISPRi) | Replogle et al,<br>2022 | RPE1 | Essential genes | 2393 | 72 |
| Jurkat-Essential<br>(CRISPRi) | Novel | Jurkat | Essential genes | 2393 | 83 |
| HepG2-Essential<br>(CRISPRi) | Novel | HepG2 | Essential genes | 2393 | 45 |
| K562-Titration<br>(CRISPRi) | Jost et al, 2020 | K562 | Essential genes | 25 genes,<br>128 guides | 143 |
| Sox9-Titration<br>(dTAG) | Naqvi et al,<br>2023 | iCNCC | SOX9 | 5 dTAG degron<br>concentrations | 7 bulk<br>samples per<br>concentration |
| Polycomb-Titration<br>(dTAG) | Weber et al,<br>2021 | mESC | Ring1b, EED<br>(Simultaneous) | 4 dTAG degron<br>concentrations | 4 bulk<br>samples per<br>concentration |

In these datasets, statistical power varies widely between perturbations due to technical features of pooled screening, including biases in sgRNA synthesis and cloning, cellular sampling noise, and variable efficiency of reverse transcription and sequencing library preparation. We illustrate the ability of TRADE to disentangle these factors from true effects with two examples from the K562-Essential dataset: knockdown of *GATA1* (**Figure 2A**) and *EIF4A3* (**Figure 2B**). These two perturbations produce very similar distributions of estimated log2FoldChange, from which it is tempting to infer that they cause similar magnitudes of transcriptome-wide changes. However, analysis with TRADE, which incorporates standard errors to estimate the variance of the true log2FoldChange, infers substantial true effect size variance for *GATA1* knockdown (transcriptome-wide impact = 0.4, corresponding to an average log_2_FC magnitude of 0.63), but negligible true effect size variance for *EIF4A3* knockdown (transcriptome-wide impact = 0.004, corresponding to an average log_2_FC magnitude of 0.06) (**Figure 2A,B**). Further examination reveals that the screen sequenced only 7 cells with *EIF4A3* knockdown (as opposed to 108 cells with *GATA1* knockdown), likely leading to large sampling variance that inflated the observed effect size distribution. This example demonstrates how TRADE can help to identify perturbations with large true transcriptome-wide effect, such as knockdown of *GATA1*, a lineage-defining transcription factor, while appropriately identifying largely null perturbations such as knockdown of *EIF4A3*, in the setting of variable power.

**Figure 2:**
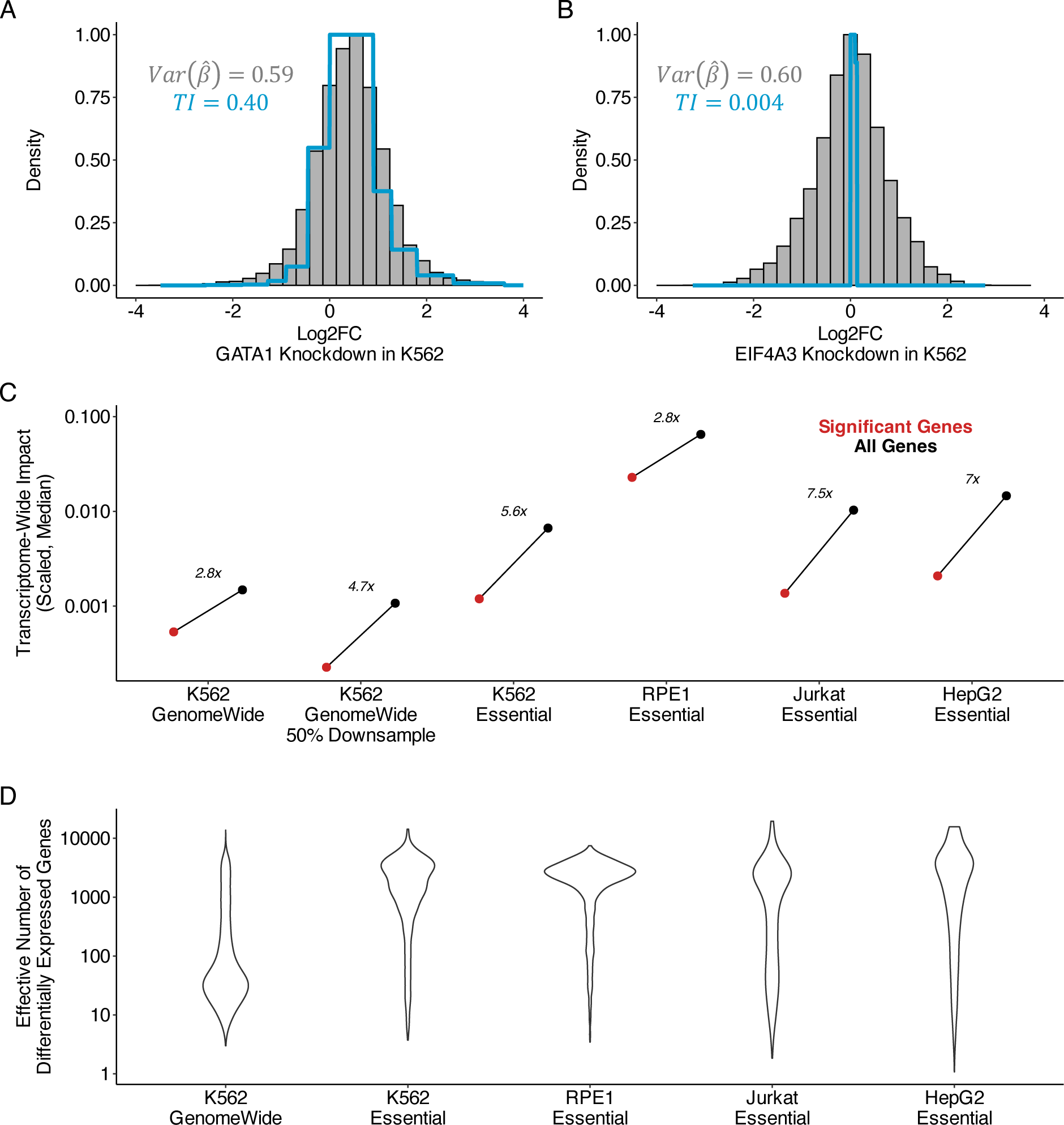
Transcriptome-wide analysis of genome-wide Perturb-Seq. **(A)** Examples of empirical log2FoldChange distribution and TRADE inferred distribution for perturbation of *GATA1* in K562 cell line. TI = transcriptome-wide impact. **(B)** Similar for perturbation of *EIF4A3* in K562. **(C)** Comparison of transcriptome-wide impact in significant and all genes in Perturb-Seq experiments. Y axis values correspond to transcriptome-wide impact estimates scaled by the number of measured genes. **(D)** Effective number of differentially expressed genes (π_*DEG*_) across Perturb-Seq datasets, for perturbations with nominally significant transcriptome-wide impact (**Methods**)

We computed the transcriptome-wide impact (see Overview of methods) of each perturbation and estimated the fraction of transcriptome-wide impact that was explained by FDR-significant effects (**Supplementary Tables 2-6**). In the K562-GenomeWide experiment, only 36% of transcriptome-wide impact was explained by FDR-significant effects (**Figure 2C**). In the four essential gene perturbation screens, we observed a similar bias where significant genes explained only a fraction of the overall transcriptome-wide impact(K562-Essential: 18%, RPE1-Essential: 35%, Jurkat-Essential: 13%, HepG2-Essential: 14%). Across all cell types, we confirmed that the transcriptome-wide impact was minimal in a negative control analysis of non-targeting guide RNAs (**Supplementary Figure 2**).

We conducted a downsampling analysis, repeating our analysis of the K562-GenomeWide experiment using only 50% of the 10x Genomics gemgroups. Whereas the signal in significant genes decreased substantially, our estimate of the total cumulative differential expression remained relatively consistent (**Figure 2C**). The small decrease in estimated transcriptome wide impact with downsampling was caused by TRADE producing conservative estimates in the setting of non-significant point estimates (**Supplementary Figure 3**). Similarly, examining perturbations which are shared between the K562-GenomeWide and K562-Essential experiments, we found that estimates of transcriptome-wide impact were far more consistent across experiments than the number of significant differentially expressed genes (transcriptome-wide impact R^2^ = 59.7%; number of DEGs R^2^ = 28.4%; **Supplementary Figure 4**). This analysis illustrates the advantages of our threshold-free approach.

Our analyses suggest that significant genes do not capture the bulk of transcriptome wide impact. How many genes are required to do so? We defined the *effective number of differentially expressed genes* (π_*DEG*_) as a function of the kurtosis of the effect size distribution, following the approach of O’Connor et al (2019). This quantity captures the evenness of differential expression across the transcriptome, without making an arbitrary distinction between zero and nearly-zero effects (**Supplementary Figure 5**). We validated our estimation procedure for π_*DEG*_ in simulations, finding that π_*DEG*_ estimates are well-calibrated, producing conservative estimates (**Supplementary Figure 6**). For the K562-GenomeWide experiment, the median π_*DEG*_was 45, suggesting that typically, tens of genes are required to explain the bulk of the transcriptome-wide impact (**Figure 2D**). Some genetic perturbations had much larger π_*DEG*_; in particular, knockdown of essential gene perturbations in all four cell types analyzed had median π_*DEG*_ greater than 500 (**Figure 2D**). In a simplified model where effects are either null or normally distributed with some variance σ^%^, π_*DEG*_ equals the number of non-null effects. Under this model, σ^%^ is equal to the ratio between the scaled transcriptome-wide impact and π_*DEG*_, and can be used to compute a typical log2FoldChange σ (**Supplementary Appendix 1**). We find that σ is largely contained in the interval [0.1,1], with subtle variation across cell type, and smaller estimates for essential versus non-essential gene perturbations (**Supplementary Appendix 1**).

### Two types of gene-set enrichment

Some sets of genes may produce greater-than-average transcriptome-wide impact when perturbed, and others may be enriched for differential expression response to perturbations of other genes. We stratified genetic perturbations by features of the targeted genes including: level of expression, effect on cellular growth (I.e. essentiality) (Meyers et al. 2017), level of selective constraint in gnomAD (Karczewski et al. 2020), and subcellular localization of their protein product (Binder et al. 2014) (**Methods**). We quantified two types of enrichment: (1) the *perturbation impact enrichment*, which captures greater-than-expected and less-than-expected transcriptome-wide impact of perturbations, and (2) the *perturbation response enrichment*, which quantifies the effect of all other perturbations on genes in the selected set (**Figure 3; Methods; Supplementary Table 7**). We focused this analysis on the K562-GenomeWide dataset, as this comprehensive dataset uniquely empowers unbiased enrichment estimation. We validated our approach with two control gene sets, one known to be enriched for perturbation response (“DE Prior”; Crow et al. 2019), and one known to be depleted of perturbation response effects (stably expressed genes; Lin et al. 2019) (**Supplementary Figure 7**). Additionally, we confirmed that genes with more efficient CRISPRi knockdown were not enriched for perturbation impact, suggesting that inter-gene variability in transcriptome-wide impact is not driven by technical factors related to CRISPRi knockdown (**Supplementary Figure 8**).

**Figure 3:**
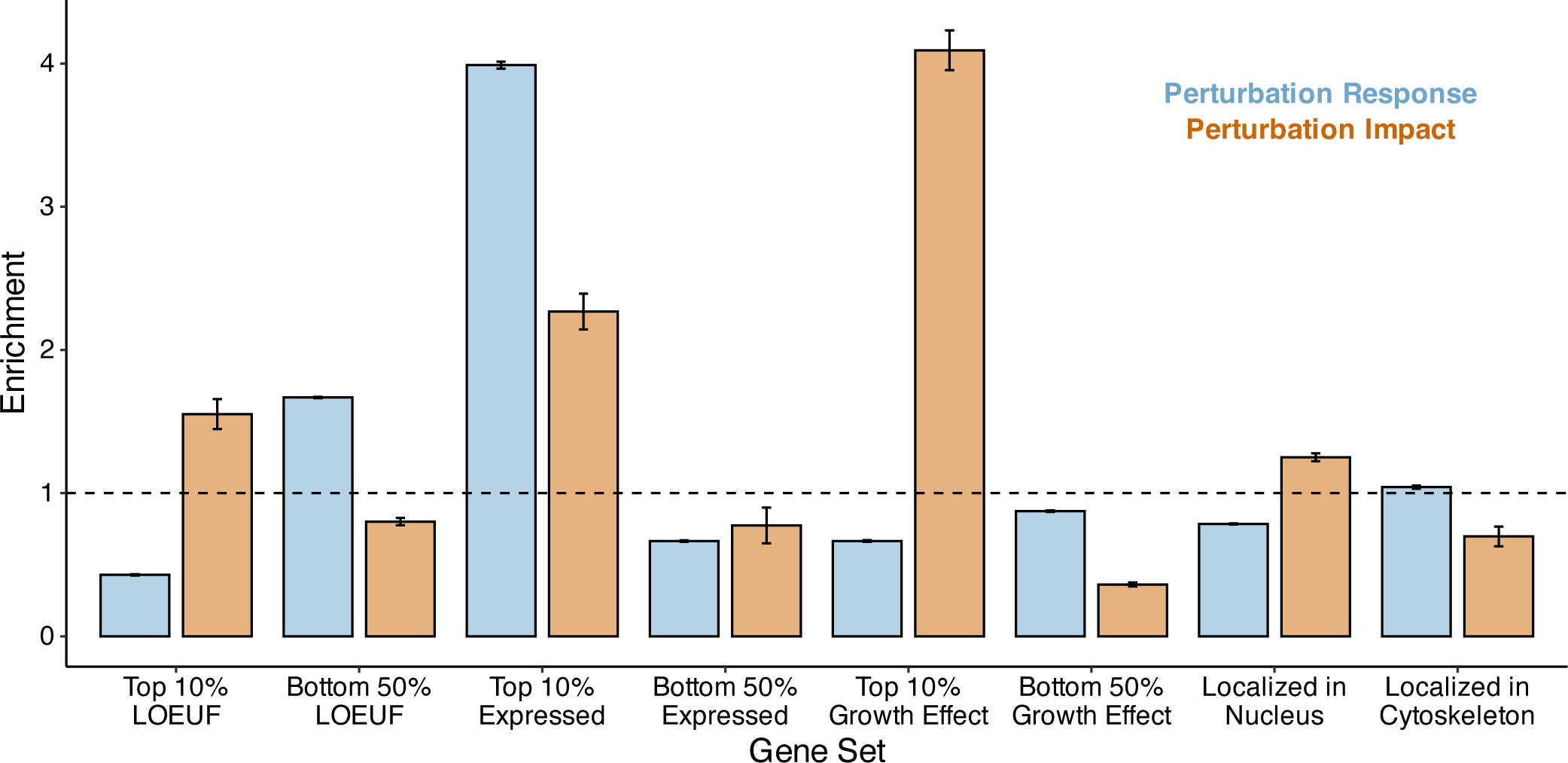
Transcriptome-wide analysis of genome-wide Perturb-Seq. TRADE-derived enrichment estimates for multiple gene sets. Blue bars represent perturbation response enrichment, the enrichment of differential expression in response to perturbations. Tan bars represent perturbation impact enrichment, the enrichment of effects on other genes when genes in that gene set are perturbed.

We found that constrained genes, which are depleted of loss-of-function variation in the general population, are enriched for perturbation impact by ∼1.57x, consistent with their functional importance. On the other hand, they are strongly depleted for perturbation response, by 0.40x, suggesting that across genes, population-level constraint is mirrored by regulatory robustness. Similarly, genes with a strong growth effect in K562 cells (roughly, those that are essential in culture) are strongly enriched for perturbation impact, by 4.22x, while being depleted for perturbation response by 0.71x. In contrast, genes that are highly expressed in K562 cells are strongly enriched for both perturbation impact (2.26x) and perturbation response (4.44x), supporting a correlation between absolute expression and functional importance. We observed only a modest perturbation impact enrichments for genes that were localized to the nucleus (1.27x), despite their direct role in transcriptional regulation; cytoskeleton-localizing genes were modestly depleted of perturbation impact (0.68x).

### Consistency of transcriptome-wide effects across cell types

The effect of perturbing a gene may vary across cell types, particularly if it participates in cell-type dependent functions. These perturbation effects may vary both in magnitude and in which genes are affected. Using data from common essential gene perturbations in the four cell lines (**Table 1**), we (1) compared transcriptome-wide impact across cell lines and (2) estimated the correlation between differential expression effects from each experiment using a bivariate extension of TRADE. We refer to this quantity as the “transcriptome-wide impact correlation” (**Methods**).

As expected, transcriptome-wide impact was correlated across cell types (average correlation = 0.62; **Supplementary Figure 8**). On average, transcriptome-wide impact was larger in the RPE1 cell line than in the other three, indicating that this cell line is more sensitive to generic perturbations than the others. A few perturbations did have greater-than-expected effects in specific cell types (**Figure 4A**). Using a liberal threshold (**Methods**), we identified 241 such perturbations (K562: 47; RPE1: 118; Jurkat: 10; HepG2: 66) (**Supplementary Table 8**). Some of these perturbations are known to be indispensable for their corresponding cell type, including *GATA1* for erythroid cells such as K562 (Weiss, Keller, and Orkin 1994) and *HMGCR* for T-cells such as Jurkat (Lacher et al. 2017), but most had no previously documented explanation for their cell-type dependent effects. As this dataset targeted primarily common essential genes which are expected to be important for growth across most cell types, there are expected to be many more examples of cell-type-specific effects in a larger cellular perturbation atlas.

**Figure 4:**
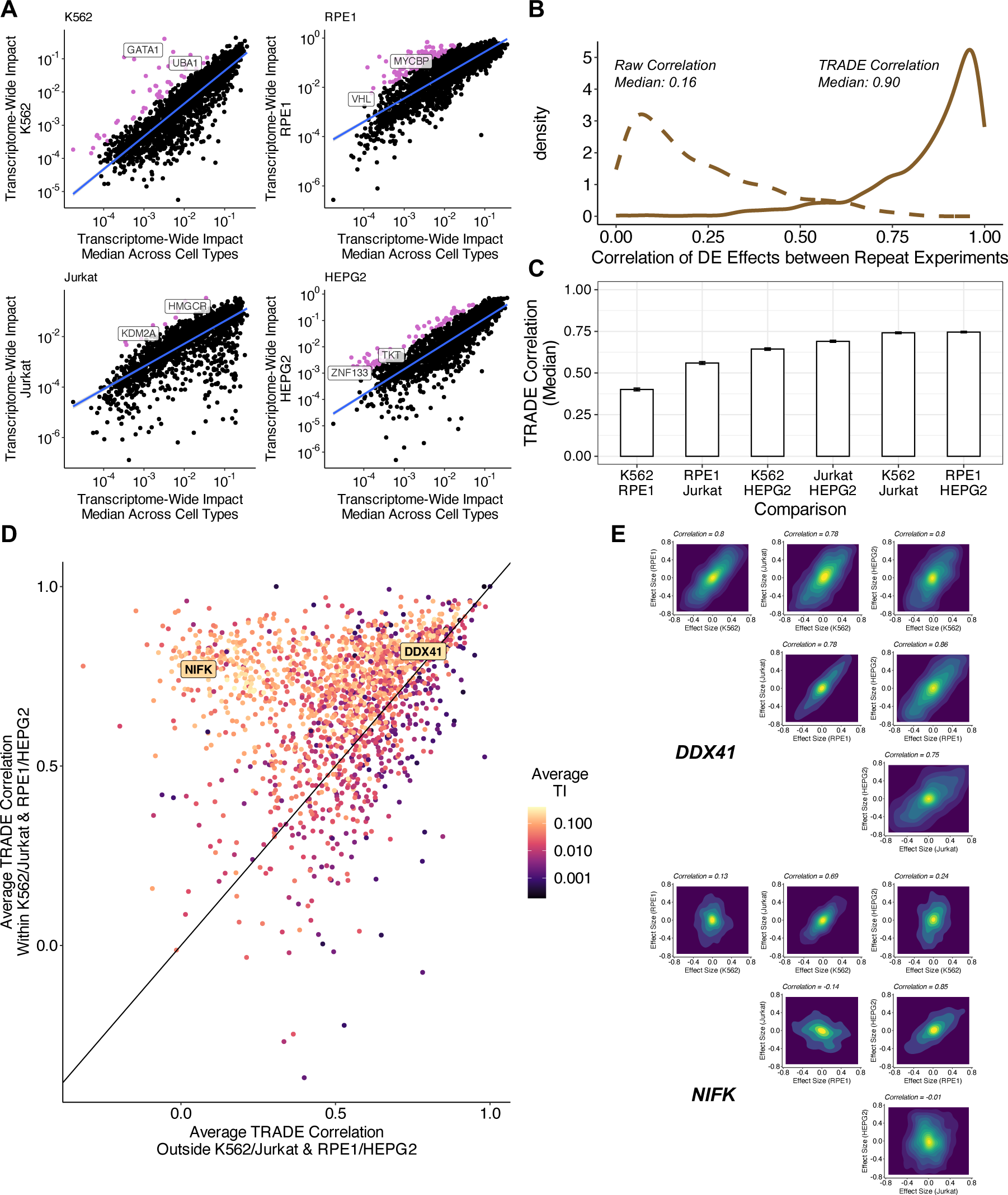
Correlation of Differential Expression Across Cell Types. **(A)** Transcriptome-wide impact of gene perturbations in each cell type versus the median across cell types, with outliers (pink) defined as being more than 1.64 standard deviations away from the fit line **(B)** Correlation of differential expression effects across replicate perturbations in K562. Dotted line represents raw correlation, solid line represents correlation estimated with TRADE. **(C)** Median correlation of perturbation effects for common essential genes for each pair of cell types. **(D)** Comparison of effect size correlation strength within similar cell types and outside of similar cell type pairs. **(E)** Examples of inferred joint effect size distributions across all pairs of cell types for perturbations of *DDX41* and *NIFK*.

Before computing correlations between different cell types, we first compared differential expression effects of the same genetic perturbations in repeated experiments (K562-GenomeWide and K562-Essential) (**Supplementary Table 9**). The median correlation between log-fold-change point estimates - not using TRADE - was only 0.16, suggesting very low replicability (**Figure 4B**). However, the median transcriptome-wide impact correlation between replicates was 0.90, implying excellent replicability (**Figure 4B**). This difference underscores the value of modeling sampling variance when estimating effect-size correlations (as uncorrelated sampling variation causes downward bias in correlation estimates; **Supplementary Appendix 2**). A few perturbations did have low between-experiment transcriptome-wide impact correlations; most of these had very low transcriptome-wide impact, and thus, their correlations are expected to be noisy (**Supplementary Figure 9**).

We used TRADE to estimate the correlation of transcriptome wide effects for perturbations of 2,053 shared essential genes across K562, RPE1, Jurkat, and HepG2 (**Supplementary Table 10**). Because these correlations are not defined in the setting of null transcriptome-wide impact, we restricted our analysis to 1660 perturbations with significant transcriptome-wide impact in all four cell types, using a very liberal threshold (Z > 0.5, corresponding to a p-value of roughly 0.3). The median transcriptome-wide impact correlation varied across pairs of cell types (**Figure 4C**). The highest median correlations were for K562/Jurkat (median correlation: 0.74) and HepG2/RPE1 (median correlation: 0.75). These functional results seem to correspond to known shared features of these cell lines: K562 and Jurkat are hematopoietic cell lines that are p53 mutant and grow in suspension, while HepG2 and RPE1 are epithelial cell lines that are p53 wild-type and are adherent. Outside of these pairs, we observed slightly weaker correlations for K562/HepG2 (0.64) and Jurkat/HepG2 (0.69), and still weaker correlations for K562/RPE1 (0.40) and RPE1/Jurkat (0.56), suggesting that RPE1 cells tend to have especially unique responses to perturbations. We considered the effect of ascertaining shared essential genes on this analysis, and determined that inferred correlations did not vary strongly with essentiality (**Supplementary Figure 10**).

Across perturbations, we observed two patterns of inter-cell-type correlations (**Figure 4D**). Some perturbations, such as knockdown of *DDX41,* had high correlations across all four cell types (**Figure 4E**). Other perturbations, such as knockdown of *NIFK,* had much higher correlations within the pairs K562/Jurkat and RPE1/HepG2 than other cell type pairs (**Figure 4E**). Clustering these perturbations with a Gaussian mixture model (**Methods**), we found that 56% of the perturbations had high correlations across all cell types (mean correlation within similar cell type pairs: 0.75; outside similar pairs: 0.66); 44% had higher correlations across similar cell types (mean correlation within similar cell type pairs: 0.61; outside similar pairs: 0.35).

### Dosage sensitivity of transcriptome-wide impact

In the experiments analyzed above, CRISPRi guide RNAs were carefully engineered to maximize on-target knockdown. Another area of significant focus in cell biology and human genetics is in generating datasets with engineered or natural variation dosage (i.e. “allelic series”) to study dosage-response relationships, which can yield insight into gene regulation and guide therapeutic design (Collins et al. 2022; Domingo et al. 2024). Traditional analytic methods struggle to compare the effects of strong to weak perturbations in these datasets as genuine response differences may be conflated with difference in signal-to-noise ratio. We reasoned that TRADE could help contend with this challenge. We applied TRADE to data from experiments that interrogated dosage-dependent transcriptome effects of depleting essential genes in K562 (Jost et al. 2020), Sox9 in induced human cranial neural crest cells (Naqvi et al. 2023), and two essential Polycomb subunits in mouse embryonic stem cells (Weber et al. 2021). Jost et al (2020) titrated gene expression with CRISPRi, which prevents transcription, and can be tuned by engineering attenuated guide RNAs containing mismatches to their target genes. Naqvi et al (2023) and Weber et al (2021) directly depleted protein levels with the dTag degron system, which can be tuned by titrating a small molecule (Nabet et al. 2018). We quantified (1) the magnitude of the transcriptome-wide impact as a function of dosage and (2) the correlation of these effects between each pair of dosages.

As expected, stronger perturbations had consistently larger transcriptome-wide impact (**Figure 5A)**. For dTAG depletion of Sox9 (**Supplementary Table 11**) and Polycomb (**Supplementary Table 13**), the transcriptome-wide impact dosage-response curve was nonlinear. Weak-to-moderate perturbations of these proteins caused relatively small transcriptome-wide effects, whereas strong perturbations caused disproportionately large transcriptome-wide effects. These genes are haploinsufficient (pLI = 1; Karczewski et al, 2020), indicating that 50% depletion is deleterious; our results suggest that stronger depletions nonetheless produce progressively larger cellular effects. We generally observed similar dose-response curves for CRISPRi knockdown of 25 essential genes in K562 cells, with varying degrees of non-linearity (**Figure 5A, Supplementary Table 15, Supplementary Figure 11)**.

**Figure 5:**
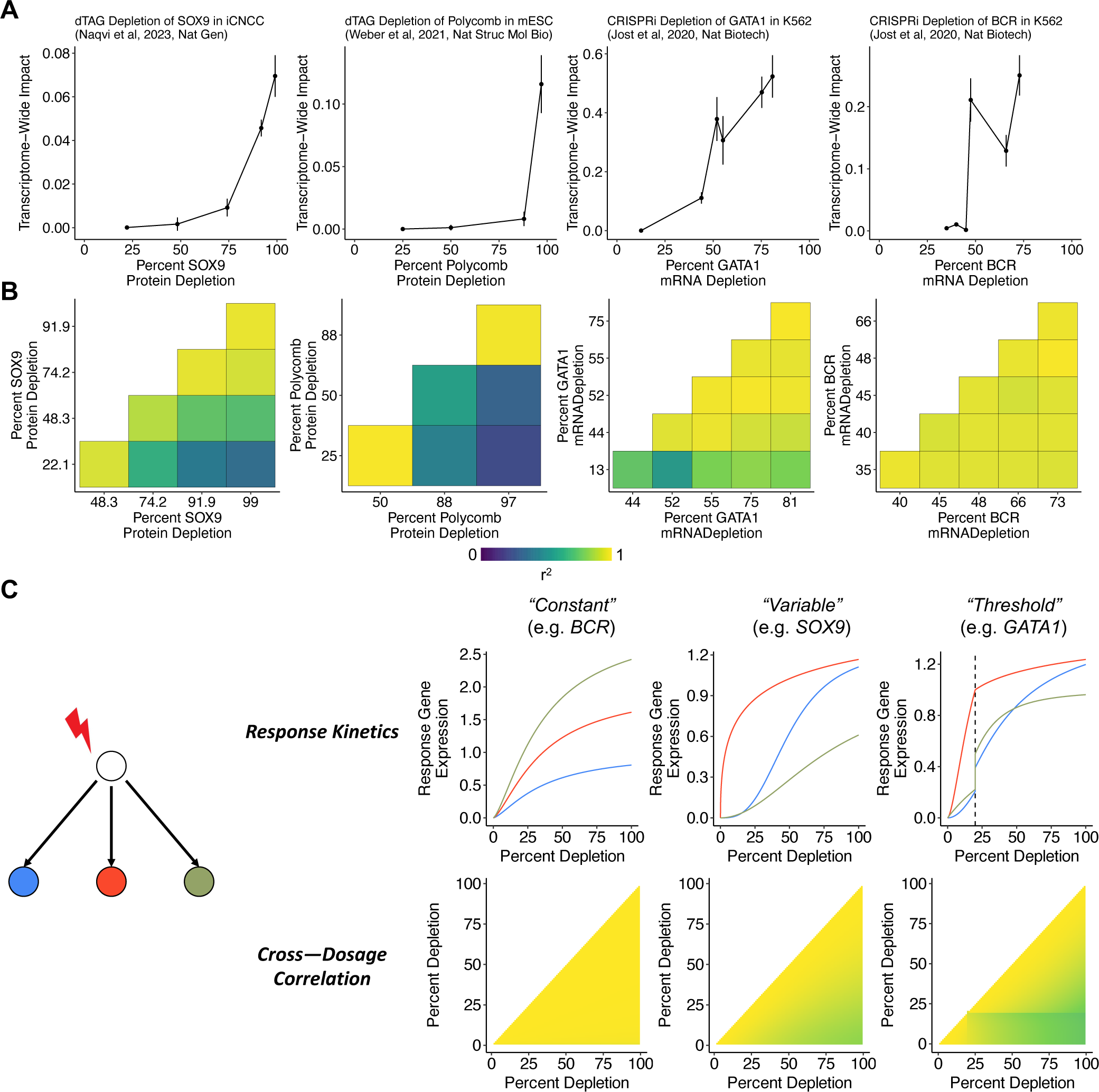
Dose-Response Relationships. **(A)** Relationship between gene dosage and transcriptome-wide impact across four experiments. **(B)** Correlations between differential expression effects at different dosages for each experiment **(C)** Observations from a toy model of perturbation effects, demonstrating relationship between response kinetics consistency and resulting pattern of cross-dosage correlations.

**Figure 6:**
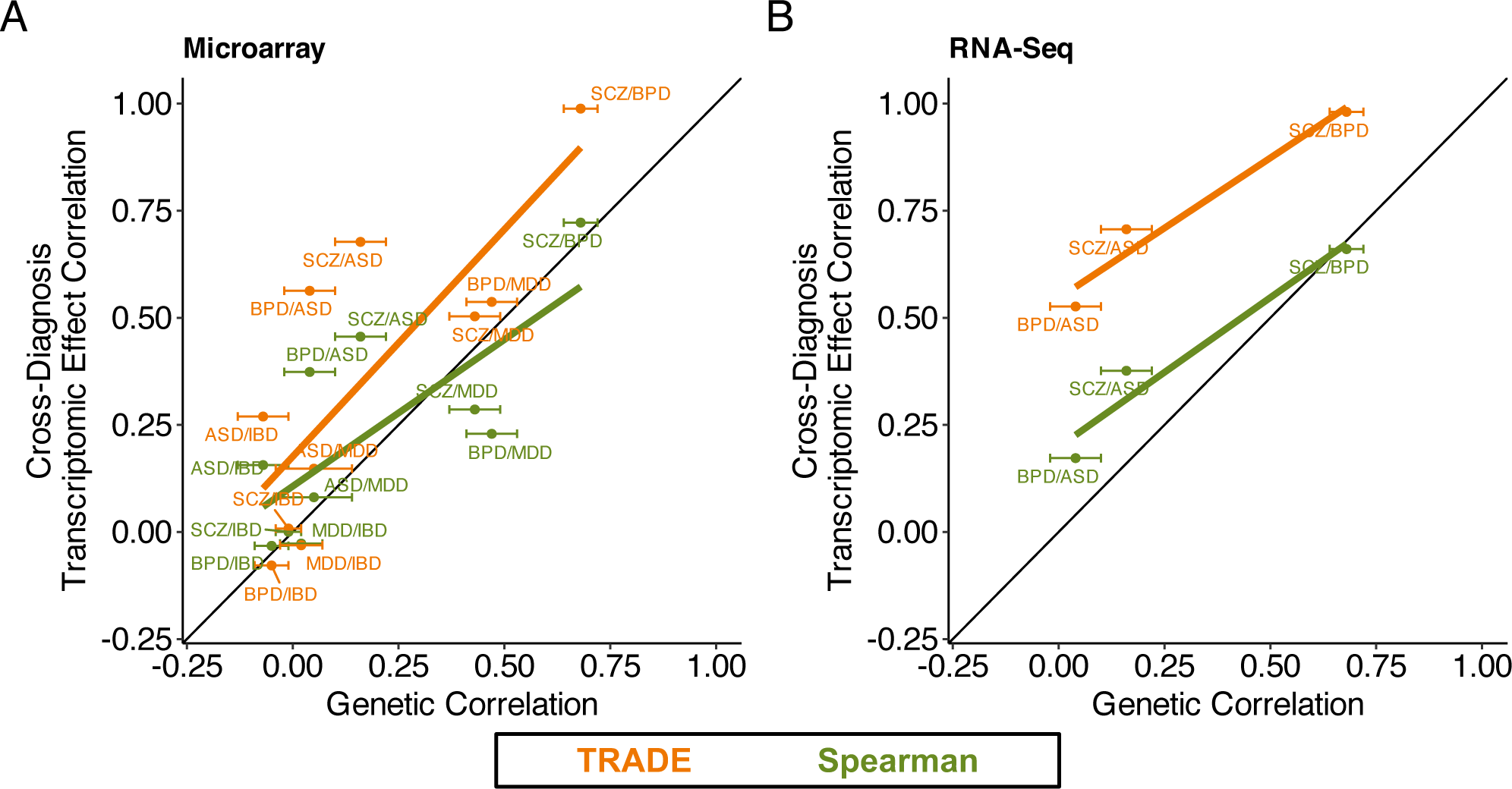
Transcriptomic Correspondence of Neuropsychiatric Conditions. Across several case/control datasets for neuropsychiatric diagnoses, estimated transcriptome-wide impact correlation (orange), compared with spearman correlations of point estimates (green). **(A)** Estimates for microarray datasets from PsychENCODE **(B)** Estimates for RNA-Seq datasets from PsychENCODE.

We next quantified the transcriptome-wide impact correlation between dosage levels for each perturbation (**Figure 5B**). For dTag depletion of Sox9 (**Supplementary Table 12**) and Polycomb (**Supplementary Table 14**), transcriptome-wide impact correlations decayed smoothly with the difference in dosage, and the smallest and largest perturbations were only moderately correlated (r=0.60, 0.48), implying that weak and strong perturbations have qualitatively different transcriptional consequences. The CRISPRi knockdown of essential genes produced a range of patterns (**Supplementary Table 16**). For example, the response to *BCR* knockdown was highly correlated across all dosage levels, despite substantial differences in the magnitude of responses (**Figure 5B**). In contrast, the response to *GATA1* knockdown was highly correlated among all but the weakest perturbation, which was only moderately correlated with the strongest perturbation (**Figure 5B**). Across the other K562 essential gene titration experiments, we found a diversity of correlation patterns, including gradient-like patterns (e.g. *ATP5E*), highly-correlated patterns (e.g. *POLR2H*), and threshold patterns (e.g. *RAN*) (**Supplementary Figure 12**).

We interpret these correlations as a readout of how the dose-response curve varies across target genes. If all downstream genes have identical response curves (up to multiplication by a constant), then the effect of a partial depletion is a fixed fraction of the effect of a full depletion, leading to a cross-dosage correlation of exactly one (**Figure 5C; Supplementary Appendix 3**). However, if the response curve varies between target genes, the correlation is less than 1, to an extent that depends on the variability of response curves (**Figure 5C; Supplementary Appendix 3**). Indeed, Naqvi et al (2023) found that a subset of Sox9 targets are sensitive at partial dosage depletions, whereas a much larger set of targets are affected only at full dosage depletions. The presence of a threshold, where the response curves change abruptly, leads to a large change in correlation magnitude across the dosage threshold. In simulations, we recapitulated the three correlation patterns described above with different sets of response curves (**Figure 5C; Methods**).

In genetic experiments and genetic association studies, it is common to study the effect of a gene by estimating a single point on its dose-response curve, potentially missing qualitatively different dosage-dependent behavior. One classical example of this phenomenon is recessivity. More generally, even haploinsufficient genes (such as *SOX9*) can have qualitatively different effects as a function of dosage. These analyses highlight the value of studying allelic series in genetic association studies, and of designing knockdown experiments at clinically relevant dosages.

### Greater transcriptomic than genetic correspondence across neuropsychiatric conditions

Gandal et al (2018) conducted a large-scale differential expression analysis of post-mortem brain tissue from individuals with neuropsychiatric conditions, comparing them with neurotypical controls. They found that differential expression effects were correlated between conditions, and that these correlations paralleled those between the genetic effects on those conditions (the genetic correlation). Because genetic effects are usually causal, this parallel was interpreted as evidence that transcriptomic overlap reflects upstream, disease-causing processes rather than confounding or downstream effects. A notable difference between the transcriptomic and genetic analyses in this study is that the genetic correlation was estimated with a REML approach that accounts for sampling variation (Lee et al. 2012), whereas the transcriptomic effect correlation was estimated as the sample Spearman correlation between differential expression point estimates, which is downwardly biased in a sample size dependent manner (**Supplementary Appendix 2).**

We reanalyzed differential expression summary statistics from this study and estimated the transcriptome-wide impact correlation between several diagnoses (**Supplementary Table 17; Figure 5A and 5B**). Integrating data from all diagnosis pairs and technologies, we found that transcriptome-wide impact correlations were substantially larger than sample Spearman correlations, with an increase for 9/9 psychiatric trait pairs. As a result, unlike the Spearman correlation estimates, the TRADE correlation estimates were larger than the between-condition genetic correlations (**Figure 5A-B)**. In contrast, TRADE appropriately estimated lower transcriptome-wide impact correlations between psychiatric diagnoses and irritable bowel disease (IBD), a non-psychiatric control trait (**Supplementary Table 17**). One explanation for this difference is that transcriptomic effects are often downstream of condition liability, and these downstream effects are often shared between neuropsychiatric conditions. Another possibility is that there exist confounding factors associated with gene expression and neuropsychiatric diagnoses in general.

One such axis of technical variation may be related to experimental assay. Studies such as PsychENCODE often integrate cohorts that profile gene expression with different technologies, such as DNA microarrays and RNA sequencing. For three conditions with independent microarray and RNA sequencing cohorts in PsychENCODE (autism, bipolar disorder, and schizophrenia), we used TRADE to estimate the correlation of transcriptomic effects between assays (**Supplementary Table 18**). The transcriptome-wide impact correlation was 0.96, 0.91, and 0.78 for autism, bipolar disorder, and schizophrenia respectively (**Supplementary Figure 13**). These estimates imply that at least in this study, most differential expression effects replicate between assays.

## Discussion

Transcriptomics is a cornerstone of modern biology. With it, questions surrounding differential expression have become ubiquitous. For many such questions, especially those that involve patterns across genes or experiments, a conventional significance-testing framework may produce misleading results. We show that these limitations can be addressed by modeling the distribution of differential expression effects explicitly via TRADE. We found that significant genes capture only a fraction of transcriptome-wide impact in large-scale Perturb-Seq experiments. Across cell types or even replicate experiments, the concordance between estimated effect sizes is attenuated due to sampling variation, but we showed that in many cases, the true effect sizes are highly concordant. In dose-response experiments, we found that dosage affects not only the magnitude of the transcriptome-wide effect, but also the genes that are affected. In a case-control as opposed to perturbational dataset, we found that the same advantages apply, and that our approach changes the interpretation of a key analysis of neuropsychiatric conditions.

The ubiquity of small differential expression effects is connected to an existing division in the field, between approaches that test for differential expression of single genes (e.g. DESeq2; Love, Huber, and Anders 2014) and those that test for differential abundance of cellular states (e.g. covarying neighborhood analysis, CNA; Reshef et al. 2022). These methods approach differential expression with distinct priors: that changes in expression will be largely restricted to a small number of genes with large effect, or that changes in expression will be spread across many hundreds or thousands of genes, reflecting a change in cell state. Estimates from TRADE, in particular π_*DEG*_, can contextualize these approaches by quantifying the degree to which differential expression is concentrated in specific target genes, versus spread across the transcriptome.

In addition to studying perturbations, an important application of differential expression analysis is to understand differences between cell types. Many analyses of cell-type variation require a *distance metric*, a scalar summary of the transcriptomic difference between groups of cells, and many such metrics have been proposed (Ji et al. 2023). Transcriptome-wide impact may be a suitable distance metric for such analyses, as it is unbiased at finite sample size (unlike the commonly used Euclidean distance, **Supplementary Appendix 4**), is easily interpretable, and can be computed from differential expression summary statistics. Indeed, we found that compared to Euclidean distance, transcriptome-wide impact produced a more coherent cell-type hierarchy of peripheral blood mononuclear cells in the OneK1K dataset (Yazar et al. 2022; Kang et al. 2023; Methods) (**Supplementary Figure 14**; **Supplementary Table 19**). However, a limitation of TRADE is that it relies upon predefined labels, and cannot be used to cluster cells into cell types.

For genetic perturbations, parameters such as transcriptome-wide impact are likely driven by the pattern of causal regulatory connections between genes, i.e. the *gene regulatory network* (GRN). Inference of GRNs from single-cell measurements is a challenging, unsolved technical problem (Pratapa et al. 2020). We speculate that, just as inferring transcriptome-wide impact is easier than inferring gene-specific effect sizes, estimating global features of the GRN may be easier than identifying individual edges. This could be achieved by pairing TRADE with a model relating the distribution of differential expression effects to GRN features such as the degree distribution or modularity. We speculate that the true GRN is densely interconnected with relatively low modularity, based on our observation that virtually all high transcriptome-wide impact perturbations also affect a large number of genes, approaching the number of genes that are expressed (**Supplementary Appendix 1**).

A key limitation of TRADE is that it currently uses only a simple readout from single cell RNA-seq experiments, the pseudo-bulk mean RNA expression level. Average expression is a widely used and highly interpretable readout, but the transcriptional state of individual cells may vary in ways that are poorly captured in pseudo-bulk (for example, due to the presence of multiple cell types) and are better understood with modeling of cell type variability (Lopez et al. 2018). In addition, some biological processes are better assayed using alternative modalities, including mRNA splicing, chromatin state, protein level, and imaging, all of which are now being studied at scale with single cell CRISPR screens (Rubin et al. 2019; Feldman et al. 2019; Gu et al. 2023; Kudo et al. 2023; Binan et al. 2023; Xu et al. 2023) We predict that future methods building on our approach will have broad application to these other phenotypic readouts as well as to the study of non-genetic perturbations such as drugs and development.

An emerging goal of functional genomics is the generation of perturbational cell atlases across multiple cellular contexts (Rood et al. 2022; Morris et al. 2024). However, as with all screening methods, there is a tradeoff where the number of assayed perturbations is ultimately constrained by experimental cost. TRADE shifts the balance in this tradeoff by allowing stable quantification of highly informative metrics including transcriptome-wide impact, correlations between perturbations, and context-dependent effects at much shallower sampling depths.

Combined with developments in screen compression (Yao et al. 2023) and cheaper sequencing technologies (Simmons et al. 2023) our method suggests a productive path toward massive scale perturbational atlases.

## Methods

### Experimental Model and Subject Details: Perturb-Seq of essential genes in Jurkat and HepG2

#### Cell line generation and maintenance

All cell lines were grown at 37°C in the presence of 5% CO2 in standard tissue culture incubators.

A CRISPRi Jurkat cell line expressing dCas9-BFP-KRAB (KOX1-derived) was obtained from the UC Berkeley Cell Culture Facility (cIGI1) and was used for growth screens. A second CRISPRi Jurkat cell line expressing the optimized UCOE-EF1α-Zim3-dCas9-P2A-mCherry CRISPRi construct was generated as previously described (Replogle *et al*., eLife 2022) and was used for Perturb-seq. Jurkat cells were grown in RPMI-1640 medium with 25 mM HEPES, 2.0 g/l NaHCO3, and 0.3 g/l L-glutamine (Gibco) supplemented with 10% (v/v) standard FBS, 2 mM glutamine, 100 units/ml penicillin, and 100 µg/ml streptomycin (Gibco).

A CRISPRi HepG2 cell line expressing UCOE-EF1α-dCas9-BFP-KRAB (KOX1-derived) was obtained from Torres *et al*. (Torres *et al*., eLife 2019), and was used for both growth screens and Perturb-seq. HepG2 cells were grown in EMEM with 1.5 g/L NaHCO3, 110 mg/L sodium pyruvate, 292 mg/L l-glutamine (Corning) supplemented with 10% (v/v) standard FBS, 100 units/mL penicillin, and 100 µg/mL streptomycin (Gibco).

HEK293T cells were used for generation of lentivirus. HEK293T cells were grown in DMEM with 25 mM d-glucose, 3.7 g/L NaHCO3, 4 mM l-glutamine (Gibco) supplemented with 10% (v/v) standard FBS, 2 mM glutamine, 100 units/ml penicillin, and 100 µg/ml streptomycin (Gibco).

#### Lentiviral production

To produce lentivirus, HEK293T cells were co-transfected with transfer plasmids and standard packaging vectors expressing VSV-G, Gag/Pol, Rev, and Tat using TransIT-LTI Transfection Reagent (Mirus). Viral supernatant was harvested 2 days after transfection and frozen at -80°C prior to transduction.

#### Library design and growth screens

Dual-sgRNA CRISPRi lentiviral libraries were previously described (Replogle *et al.,* Cell 2022). Briefly, a preliminary sgRNA library (dJR058, n=2291 dual-sgRNA elements) with even representation of all dual-sgRNA constructs was used for growth screens. This library contains a single dual-sgRNA construct targeting i) 20Q1 Cancer Dependency Map common essential genes (https://depmap.org/portal/download/) and (ii) 5% non-targeting control sgRNAs cloned into pJR101 (Addgene #187241). A second sgRNA library (dJR092, n=2688 dual-sgRNA elements, Supplementary Table 20) which adjusted the representation of sgRNAs to decrease dropout of essential genes was used for Perturb-seq experiments. The sgRNA abundance was corrected according to the effects observed in growth screens (described below); for example, a guide with roughly four-fold depletion in growth screens was four-fold overrepresented in dJR092. This library also included sgRNAs targeting a number of additional genes with interesting phenotypes in the K562 genome-wide Perturb-seq dataset.

Pooled growth screens in Jurkat cells were performed by transducing Jurkat cells expressing dCas9-BFP-KRAB (cIGI1) with the dual-sgRNA CRISPRi lentiviral library, dJR058. Screens were performed in biological replicate maintaining a coverage of >1000 cells per library element for the duration of the screen. Cells were transduced by spinfection (1000g) with polybrene (8 µg/mL, Sigma-Alrich) to obtain an infection rate of 10%-20%. On day 3 post-transduction, cells were sorted to near-purity by FACS (FACSAria2, BD Biosciences), using GFP as a marker for sgRNA vector transduction. On day 7 post-transduction, an aliquot of cells was harvested for sequencing to compare sgRNA abundances to the plasmid library.

Pooled growth screens in HepG2 cells were performed by transducing HepG2 cells expressing dCas9-BFP-KRAB with the dual-sgRNA CRISPRi lentiviral library, dJR058. Screens were performed in biological replicate maintaining a coverage of >1000 cells per library element for the duration of the screen. Cells were transduced by plating in viral supernatant with polybrene (8 µg/mL, Sigma-Alrich) to obtain an infection rate of 30%-40% based on GFP measurement by FACS (FACSAria2, BD Biosciences). On day 7 post-transduction, an aliquot of cells was harvested for sequencing to compare sgRNA abundances to the plasmid library.

Library preparation and sequencing of growth screens followed the protocol previously described (Replogle *et al*., Cell 2022).

#### Perturb-seq experiments, library preparation, and sequencing

For the Jurkat Perturb-seq experiment, Jurkat cells expressing Zim3-dCas9-P2A-mCherry were transduced with dJR092 library lentivirus by spinfection (1000g) with polybrene (8 µg/mL, Sigma-Alrich) with a targeted low infection rate of ∼10%. This low rate was chosen to reduce the chances of a single cell being infected by multiple viruses. Cells were maintained at a coverage of >1000 cells per library element for the duration of the screen. On day 3 post-transduction, an infection rate of 7% was measured based on GFP as a marker of transduction, and cells were sorted to near purity by FACS (FACSAria2, BD Biosciences). On day 7 post-transduction, cells were measured to be >90% GFP positive (Attune NxT, ThermoFisher) and were prepared for single-cell RNA-sequencing by resuspension in 1X PBS with 0.04% BSA as detailed in the 10x Genomics Single Cell Protocols Cell Preparation Guide (10x Genomics, CG00053 Rev C). Cells were separated into droplet emulsions using the Chromium Controller (10x Genomics) with Chromium Single-Cell 3′ Gel Beads v3 (10x Genomics, PN-1000075 and PN-1000153) across 56 GEM groups following the 10x Genomics Chromium Single Cell 3ʹ Reagent Kits v3 User Guide with Feature Barcode technology for CRISPR Screening (CG000184 Rev C) with the goal of recovering ∼15,000 cells per GEM group before filtering.

For the HepG2 Perturb-seq experiment, HepG2 cells expressing dCas9-BFP-KRAB were transduced with dJR092 library lentiviral supernatant with polybrene (8 µg/mL, Sigma-Alrich) with a targeted low infection rate of ∼10%. Cells were maintained at a coverage of >1000 cells per library element for the duration of the screen. On day 3 post-transduction, an infection rate of 7% was measured based on GFP as a marker of transduction, and cells were sorted to near purity by FACS (FACSAria2, BD Biosciences). On day 7 post-transduction, cells were dissociated using Accutase (StemCell Technologies) for 30 minutes and resuspended in 5 mM EDTA-PBS. In order to decrease cell doublets in the single-cell RNA-sequencing, GFP positive singlets were isolated by FACS (FACSAria2, BD Biosciences) with a final cell population measured to be ∼90% GFP positive. Cells were then prepared for single-cell RNA-sequencing by resuspension in 1X PBS with 0.04% BSA as detailed in the 10x Genomics Single Cell Protocols Cell Preparation Guide (10x Genomics, CG00053 Rev C). Cells were separated into droplet emulsions using the Chromium Controller (10x Genomics) with Chromium Single-Cell 3′ Gel Beads v3 (10x Genomics, PN-1000075 and PN-1000153) across 56 GEM groups following the 10x Genomics Chromium Single Cell 3ʹ Reagent Kits v3 User Guide with Feature Barcode technology for CRISPR Screening (CG000184 Rev C) with the goal of recovering ∼15,000 cells per GEM group before filtering.

For library preparation, samples were processed according to 10x Genomics Chromium Single Cell 3ʹ Reagent Kits v3 User Guide with Feature Barcode technology for CRISPR Screening (CG000184 Rev C) with magnetic selections conducted on an Alpaqua Catalyst 96 plate (#A000550). For sequencing, mRNA and sgRNA libraries were pooled to avoid index collisions and sequenced on a NovaSeq 6000 (Illumina) according to the 10x Genomics User Guide.

### Quantification and Statistical Analysis

#### Perturb-seq in Jurkat and HepG2: alignment, cell calling, sgRNA assignment, and cell filtering

As previously described (Replogle *et al*., Cell 2022), Cell Ranger 4.0.0 software (10x Genomics) was used for scRNA-seq and sgRNA alignment, collapsing reads to UMI counts, and cell calling. The 10x Genomics GRCh38 version 2020-A genome build was used as a reference transcriptome. For sgRNA assignment, reads were first downsampled by GEM group to produce a more even distribution of the number of reads per cell, with an upper threshold of 3000 mean mapped reads per cell in the Jurkat experiment and 2500 mean mapped reads per cell in the HepG2 experiment. Guide calling was performed with a Poisson-Gaussian mixture model as previously described (Replogle *et al*., Nature Biotech 2020), with only cells bearing two sgRNAs targeting the same gene or a single sgRNA used for downstream analysis. Cells were filtered for quality to remove cells with low UMI content (Jurkat: < 14%, HepG2: <18%) and high mitochondrial RNA content (Jurkat: > 1750 UMIs, HepG2: > 3000 UMIs). These filters removed 7408 cells from the Jurkat-Essential experiment (262956 cells retained), and 15952 cells from the HepG2-Essential experiment (145473 cells retained).

A similar procedure was performed as previously described for the K562-GenomeWide, K562-Essential, and RPE1-Essential datasets (Replogle et al, Cell, 2022)

#### Modeling transcriptome-wide responses

In a differential expression experiment, read depth is quantified for each gene and each cell or sample. The resulting counts are typically modeled as following a distribution (e.g., negative binomial) whose mean may differ between two conditions, and the difference is quantified as a *log-fold change*, defined as the difference between the logarithm of population expression means between the two conditions. In a typical experiment, the log-fold change can be interpreted as the effect size of a condition or perturbation on a gene, and many biological questions are related to the distribution of true effect sizes across genes. However, we only observe the distribution of estimated effect sizes. These estimates can be modeled as the sum of two distributions: the distribution of true effect sizes, and the sampling distribution in the experiment. TRADE is a method to disentangle these components.

To do so, TRADE uses *ash* <u>(Stephens 2016)</u> to estimate the effect size distribution from differential expression summary statistics, which we compute with *DESeq2* <u>(Love et al. 2014)</u>. *DESeq2* fits a regularized negative binomial generalized linear model, sharing information across genes to improve overdispersion estimates. It has been applied to many different contrasts, including genetic perturbations and different cell types. *ash* models an effect size distribution by learning the weights of a flexible mixture model using maximum likelihood. *ash* incorporates standard error estimates into the inference procedure, effectively down-weighting noisier log-fold change estimates, such as those for very lowly expressed genes. Whereas *ash* was initially designed to estimate effect size distributions as an intermediate step prior to shrinkage, TRADE uses it to estimate the effect size distribution itself.

#### Features of the effect size distribution

The transcriptome-wide impact is defined as the variance of the distribution of differential expression effect sizes, in units of log2-fold change. The transcriptome-wide impact captures the overall degree of transcriptomic change across a contrast of interest, e.g. a perturbation.

Importantly, transcriptome-wide impact is in interpretable units of Log_2_FC^2^. For example, if a hypothetical perturbation affects all genes with normally distributed effect sizes, and the transcriptome-wide impact is 0.25, it means that a typical gene has an effect size of 0.5.

Beyond this simplistic model, a large transcriptome-wide impact may arise either because a perturbation has a large effect on a few genes, or because it has smaller effects on many of them. To distinguish between these possibilities, we define the *effective number of differentially expressed genes (*π_*DEG*_) as:

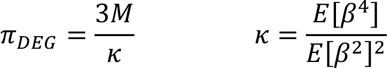

Where *M* is the number of genes with measured expression, and*k* is the kurtosis (normalized fourth moment) of the inferred effect size distribution. If a perturbation has a large effect on only a few genes, then*k* is large, and π_*DEG*_is small. Conversely, if a perturbation affects all genes with normally distributed effect sizes, then*k* equals 3, and π_*DEG*_ is equal to the number of genes (**Supplementary Figure 5**).

Different sets of genes may be enriched or depleted for differential expression. We define the perturbation response enrichment of a gene set as:

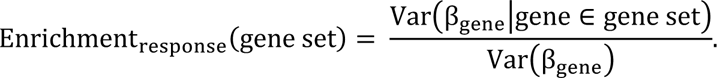

We estimate the numerator by applying *ash* to genes in the gene set, and we estimate the denominator by applying it to all genes. This approach is expected to be approximately unbiased for most gene sets. However, we do also use it to estimate the fraction of signal in FDR-significant genes; we note that such estimates are expected to be upwardly biased by winner’s curse. In Figure 3, we report the mean perturbation response enrichment across perturbations.

In addition to the perturbation response enrichment, we also estimate the perturbation impact enrichment. If *TI_i_* is the transcriptome-wide impact of perturbation *i*, the perturbation impact enrichment is:

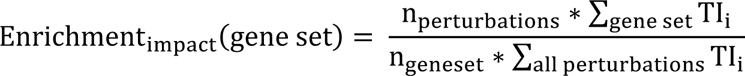

We estimate this quantity by substituting the estimated transcriptome-wide impact for the true transcriptome-wide impact.

#### TRADE Implementation details (univariate)

Briefly, to model the effect size distribution, we used *ash* to fit the following mixture model:

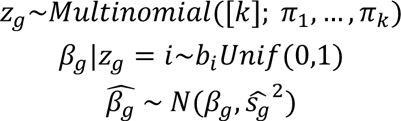

Where 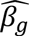 is the estimated log_2_FoldChange, β*_g_* is the true log_2_FoldChange, 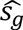 is the estimated standard error, *z*_*g*_ matches gene *g* to a mixture component uniform distribution *i* with one extremum at 0 and the other at *bi*, and π*i* are the weights for components of the mixture distribution.

*ash* fits this model with interior point optimization methods (for details, see Stephens et al, 2016, *Biostatistics*). For mixture components, we used a fine grid of uniform distributions with one extremum at zero as mixture components (“half-uniform”) rather than zero-centered uniform mixture components, to allow for estimation of asymmetric effect size distributions; Stephens (2016) found that these this model is sufficiently flexible to model realistic effect size distributions.

We largely used *ash* with default settings; we made two modifications

1. We restricted the range of mixture components to the smallest and largest observed effect size, rather than the default behavior of c(-Inf, Inf), to improve computational efficiency
2. We used a uniform rather than null-biased prior; using a null-biased prior is crucial for accurately computing the local false sign rate, but is less important for estimating the effect size distribution itself. We removed the null-biased prior in order to prevent bias in our distribution estimate.

#### Simulations

To assess the performance of TRADE, we first simulated gene expression counts for two conditions. We first used DESeq2 to estimate the expression mean and dispersion for each of the first 10 batches of control cells (i.e. cells with a non-targeting guide RNA) of the K562-GenomeWide dataset. We simulated “control” counts by sampling from negative binomial distributions with these empirical mean and dispersion estimates, for each batch. We then simulated “perturbed” counts by sampling from negative binomial distributions with “perturbed” means (i.e. multiplied by the fold change, see below) and the same dispersion, for each batch. In summary, this procedure produces a single cell expression dataset with realistic means, dispersions, and batch structure.

We then analyzed this simulated data with DESeq2 and TRADE. We generated a pseudobulk dataset by summing counts for each gene, for each condition, for each batch, creating a dataset with the number of samples equal to twice the number of batches. We then used DESeq2 to fit the following model:

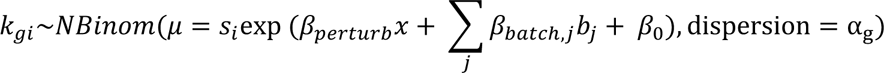

xWhere *k_gi_* is the observed expression count for gene *g* for pseudobulk observation *i*,*s_i_* is a per-observation normalization factor computed with the default DESeq2 median-of-ratios approach, *x* is a binary variable denoting the presence of a perturbation, *b_j_* is a binary variable denoting whether the pseudobulk observation comes from batch *j*, and α_*g*_ is the supra-Poisson overdispersion.

For details on fitting this model, see Love et al (2014).

To characterize estimation of transcriptome-wide impact, we generated effect sizes from 30 distinct effect size distributions:

● 3 levels of sparsity: A point-normal distribution with 95% of effects equal to zero and the other 5% drawn from a normal distribution, a point-normal distribution with 75% of effects equal to zero and the other 25% drawn from a normal distribution, and a fully infinitesimal model with 100% of effects drawn from a normal distribution
● 10 values of transcriptome-wide impact: Values ranging from 0.05 to 0.5; 0.5 is roughly the estimated value for the largest perturbations from the Replogle et al (2022) dataset. For the point normal distributions, the variance of the normal component was scaled up to equalize transcriptome-wide impact with the infinitesimal simulation (i.e. multiplying by sqrt(20) for the 95% sparse distribution and 2 for the 75% sparse distribution

To characterize estimation of π_*DEG*_, we generated effect sizes from 10 distinct effect size distributions, reflecting 10 levels of sparsity. Effect sizes were sampled from a point normal distribution with sparsity ranging from 0.05 to 0.95, with the normal component having variance 0.25

We repeated these simulations at three different sample sizes: N = 20 cells per condition, N = 200 cells per condition, and N = 2000 cells per condition. N = 200 is similar to the typical sample size regime for the Replogle et al (2022) dataset.

For each combination of parameters, we ran 100 replicate simulations.

#### Genome-wide Perturb Seq

We analyzed data from five large-scale Perturb-Seq experiments, including three from Replogle et al. 2022 (K562-GenomeWide, K562-Essential, RPE1-Essential) and two that are new (Jurkat-Essential, HepG2-Essential) (see Data Availability). We generated differential expression summary statistics (i.e. log2FoldChanges and standard errors) for each perturbation as follows:

1. We generated a per-batch (“gem-group”) pseudobulk dataset, summing counts across control cells (i.e. cells carrying a non-targeting guide RNA) and perturbed cells (i.e. cells carrying a guide RNA against a particular gene) within each batch.
2. We estimated differential expression effects from this pseudobulk dataset using DESeq2, with an identical model as in our simulations (see above)

We computed p-values with the Likelihood Ratio Test as implemented in DESeq2.

We modeled each batch as a fixed effect, and DESeq2 scales poorly with the number of covariates. This presented serious challenges only for the K562-GenomeWide dataset, which had 272 batches. To circumvent this issue, we analyzed the K562-GenomeWide dataset in four “mega-batches” of 68 batches each, and then meta-analyzed the resulting four sets of log2FoldChange estimates using inverse variance weighted meta-analysis.

#### Gene annotations

For our enrichment analyses in the K562-GenomeWide dataset, we used the following gene annotations (see Data Availability):

● Expression level: Estimated from the K562-GenomeWide dataset itself as the mean expression level.
● Growth effect: We downloaded growth effect estimates from the K562 CRISPR growth screen in the Cancer DepMap project
● Loss of Function Observed over Expected Upper Fraction (LOEUF): We downloaded these estimates from the gnomad v2 resource (Karczewski et al. 2020)
● Nuclear and cytoskeletal localization: We downloaded cellular localization annotations from the COMPARTMENTS database (Binder et al. 2014)
● DE Prior: We downloaded the DE Prior ranked list from the supplementary information of Crow et al (2019)
● Stably expressed genes (SEG): We downloaded the list of human SEGs from Lin et al (2019)
● On-Target Knockdown: We estimated the log2FoldChange for the target gene in each experiment with *DESeq2*

For the quantitative annotations, we generated two annotations, Top 10% and Bottom 50%, for enrichment analyses.

#### Transcriptome-wide analysis of Differential Expression (Bivariate)

Given two sets of differential expression summary statistics (e.g. log2FoldChanges and standard errors computed with DESeq2), we estimated the joint distribution of effect sizes using *mash* (Urbut et al, NG). *mash* fits a mixture of multivariate normal distributions to model the joint distribution of effect sizes across an arbitrary number of experiments, for example eQTLs from tens of tissues; we used *mash* to model bivariate effect size distributions, with a particular interest in estimating the correlation of effects between two perturbations.

Briefly, *mash* finds the weights π that maximize the following likelihood:

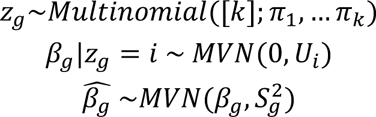

where 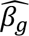 is a two-element vector of estimated effect sizes, β*_g_* is a two-element vector of true effect sizes, 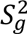 is the sampling covariance matrix of the true effect sizes, *z* matches gene *g* to a mixture component multivariate normal distribution parameterized by fixed covariance matrix *U_i_*, and π*_i_* is the weight for the component *i* of the mixture distribution

We choose *S_g_* to be a diagonal matrix with diagonal entries equal to the variance of the individual estimated effects. This choice is appropriate when each estimate is derived from a different experiment, which is the case in our analyses.

Selecting the covariance matrices *U_i_* is a crucial step in this analysis. By default, *mash* recommends a combination of “canonical” (i.e. reflecting simple correlation patterns) and data-derived (i.e. from factorization of the observed data matrix) covariance matrices, across a range of scaling factors. We used these *mash* default covariance matrices, and added several more matrices comprising an “adaptive grid”. We did so because while *mash* was designed primarily for multivariate experiments with several conditions, where specifying all possible covariance patterns is not feasible, we are interested in the bivariate case, where doing so is feasible.

We obtain this adaptive grid of covariance matrices by first running univariate *ash* in each condition, with half-normal mixture components. We then retain the component variances with non-zero weight for each distribution. Then, for each combination of variances, we create covariances matrices with several covariance values corresponding to a grid of correlations between -1 and 1 (in our experiments, 21 correlation values was a sufficiently dense grid). This procedure produces a set of covariance matrices that attempt to tile all possible bivariate relationships between the two perturbations.

#### Identification of genes with cell-type-specific perturbation effects

To identify perturbations with exceptionally large transcriptome-wide impact in one cell type, we regressed log-transformed transcriptome-wide impact estimates from each cell type on the median log-transformed transcriptome-wide impact across all four cell types. This regression included 2050 perturbations, excluding three common essential genes that had zero transcriptome-wide impact in at least one cell type. We then defined perturbations with cell-type specific effects as perturbations with a standardized residual from this regression greater than 1.64, i.e. corresponding to a p-value of 0.1.

Notably, this regression included fitted parameters for both intercept and slope, meaning that cell-type-specific effects were not identified only because one cell type exhibits stronger effects overall.

#### Clustering genetic perturbations across cell types

Visualization of the relationship between transcriptome-wide impact correlation within and between each pair of more-similar cell types (**Figure 4D**) motivated us to cluster perturbations based on these values with a bivariate gaussian mixture model. We fit a bivariate gaussian mixture model using an expectation-maximization algorithm as implemented in the *mclust* package in R. The resulting mixture components reflected the visually apparent clusters from **Figure 4D**, i.e. including one component with relatively high correlations within and outside of similar cell types (mean correlation within = 0.75, mean correlation outside = 0.66) and one component with lower correlations outside of similar cell types (mean correlation within = 0.61, mean correlation outside = 0.35). We assigned each genetic perturbation to one of the two components based on the posterior probabilities from this model.

#### Perturb-Seq with Attenuated Guide RNAs

We downloaded publicly available processed scRNA-seq data from Jost et al (2020). Full details are available in the primary manuscript describing this dataset. This data is largely identical to those described above from Replogle et al (2022), with multiple guides (with several targeting each of 25 essential genes) arrayed across three batches.

To analyze this dataset, we used an identical approach as the genome-wide and essential-wide experiments from Replogle et al (2022), performing a batch pseudobulk analysis with DESeq2. To harmonize this analysis with that of Replogle et al (2022), we limited the measured genes analyzed to those with an average expression level of 0.01 UMIs across cells. To estimate standard errors, we used a block-jackknife across cells with 100 blocks.

We estimated the degree of on-target knockdown using the Log2FoldChange for the target gene from DESeq2.

#### dTAG Depletion of SOX9

Gene-wise RNA counts were downloaded from the Zenodo archive accompanying Naqvi et al (2023), and differential expression analysis was conducted using the script from the same repository. Briefly, RNA was sequenced from bulk samples of human embryonic stem-cell derived human neural crest cells with varying concentrations of dTAG targeting Sox9. RNA-seq data was aligned with Salmon, and differential expression analysis was carried out with DESeq2, with differentiation batch as a covariate. Standard errors for the TRADE analysis were computed via a sample jackknife.

#### dTAG Depletion of Polycomb Repressive Complex

Gene-wise RNA counts were downloaded from the GEO repository accompanying Weber et al (2021). Briefly, RNA was sequenced from bulk samples of mouse embryonic stem cells with varying concentrations of dTag targeting Ring1b and Eed. RNA-seq data was aligned with kallisto, and differential expression analysis was carried out with DESeq2. Standard errors for the TRADE analysis were computed via a sample jackknife.

#### Simulations of dose-response curves

To simulate correlation of perturbation effects across dosage levels, we simulated 10000 target gene expression profiles downstream of a perturbed gene, The response function of each gene was simulated with a Hill Equation:

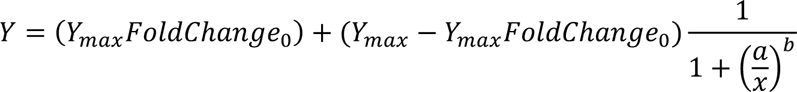

Where *Y* is the expression level of the target gene at dosage level *x* of the perturbed gene, *Y_max_* is the expression level of the target gene at full dosage of the perturbed gene, *FoldChange*_0_ is the fold change of the target gene associated with full depletion of the perturbed gene, *a* is the concentration associated with half-maximal response, and *b* is the “Hill coefficient” or the degree of cooperativity.

For all simulations, across genes, *Y*_F@G_ was drawn from the normal distribution *N*(100,5), and *FoldChange*_0_ was drawn from the normal distribution *N*(0,2) (i.e. infinitesimal architecture).

For the “Constant” simulation, *a* and *b* were constant (50 and 0.5, respectively). For the “Variable” simulation, *a* was drawn from the uniform distribution *Unif*(10,90) and *b* was drawn from the uniform distribution *Unif*(0.1,5). For the threshold simulation, *a* and *b* were drawn from these uniform distributions two times independently, to compute curves before and after a threshold of 20.

From the simulated response kinetic curves, log2FoldChanges were computed, and correlated across dosage levels.

#### Case/control differential expression in neuropsychiatric disorders

We downloaded case/control, RNA-Seq and microarray-based, differential expression summary statistics for the PsychENCODE dataset from Gandal et al (2018) for autism, schizophrenia, bipolar disorder, major depressive disorder, and irritable bowel disease. Following Gandal et al (2018), we used the estimates of cross-disorder genetic correlation from Lee et al (2013)

#### Estimating cell type hierarchies in the OneK1K dataset

We downloaded count-based sequencing data from the OneK1K cohort (Yazar et al, 2022), using the post-publication quality control of this dataset by Rumker et al (2023). We excluded one individual who had cells present in multiple batches. We generated a pseudobulk dataset by summing the counts of each individual, for each of the 28 PBMC cell types (i.e. excluding erythrocytes, hematopoietic stem and progenitor cells, and platelets) identified by Rumker et al (2023). We generated a pseudobulk dataset by summing counts within each individual, for each cell type. We then used DESeq2 to estimate differential expression between each pair of cell types, with batch as a covariate. We then used TRADE to estimate the transcriptome-wide impact between pairs of cell types.

For the Euclidean distance analysis, we took a similar to the above, but fit a DESeq2 model with only an intercept term to the pseudobulk data from each cell type, to estimate the mean expression of each gene. For each cell type, we then normalized these mean expression profiles by converting to “counts-per-10k” units (cp10k), adding 1, and log-transforming. We then computed the Euclidean distance between each pair of normalized mean cell type expression profiles.

## Code Availability

The TRADE method, with accompanying documentation, is publicly available as an R package at https://github.com/ajaynadig/TRADE.

## Data Availability

Raw sequencing data are deposited on SRA under BioProject PRJNA1100571. Aligned sequencing data and processed single-cell populations are available on GEO at GSE264667.

## Acknowledgements

AN is supported by NIH grant F31HG013036. JMR is supported by NIH grants F31NS115380 and T32GM007618. This work was funded by the National Institutes of Health (NIH) Center of Excellences in Genome Sciences (JSW). The project described was supported by award Number T32GM007753 and T32GM144273 from the National Institute of General Medical Sciences. The content is solely the responsibility of the authors and does not necessarily represent the official views of the National Institute of General Medical Sciences or the National Institutes of Health. JSW is an HHMI investigator. We thank K.A. Lagattuta, D.J. Weiner, B. Harris, T. Aicher, D.L. Barabasi, K. Maher, T. Kamath, M.T. Tegtmeyer, and members of the O’Connor and Robinson labs for helpful comments and discussions.

## References

Adamson, Britt, Thomas M. Norman, Marco Jost, Min Y. Cho, James K. Nuñez, Yuwen Chen, Jacqueline E. Villalta, et al. 2016. “A Multiplexed Single-Cell CRISPR Screening Platform Enables Systematic Dissection of the Unfolded Protein Response.” Cell 167 (7): 1867–82.e21.

Binan, Loϊc, Serwah Danquah, Vera Valakh, Brooke Simonton, Jon Bezney, Ralda Nehme, Brian Cleary, and Samouil L. Farhi. 2023. “Simultaneous CRISPR Screening and Spatial Transcriptomics Reveals Intracellular, Intercellular, and Functional Transcriptional Circuits.” bioRxiv : The Preprint Server for Biology, December. 10.1101/2023.11.30.569494.

Binder, Janos X., Sune Pletscher-Frankild, Kalliopi Tsafou, Christian Stolte, Seán I. O’Donoghue, Reinhard Schneider, and Lars Juhl Jensen. 2014. “COMPARTMENTS: Unification and Visualization of Protein Subcellular Localization Evidence.” Database: The Journal of Biological Databases and Curation 2014 (February): bau012.

Bulik-Sullivan, Brendan, Hilary K. Finucane, Verneri Anttila, Alexander Gusev, Felix R. Day, Po-Ru Loh, ReproGen Consortium, et al. 2015. “An Atlas of Genetic Correlations across Human Diseases and Traits.” Nature Genetics 47 (11): 1236–41.

Collins, Ryan L., Joseph T. Glessner, Eleonora Porcu, Maarja Lepamets, Rhonda Brandon, Christopher Lauricella, Lide Han, et al. 2022. “A Cross-Disorder Dosage Sensitivity Map of the Human Genome.” Cell 185 (16): 3041–55.e25.

Crow, Megan, Nathaniel Lim, Sara Ballouz, Paul Pavlidis, and Jesse Gillis. 2019. “Predictability of Human Differential Gene Expression.” Proceedings of the National Academy of Sciences of the United States of America 116 (13): 6491–6500.

Dixit, Atray, Oren Parnas, Biyu Li, Jenny Chen, Charles P. Fulco, Livnat Jerby-Arnon, Nemanja D. Marjanovic, et al. 2016. “Perturb-Seq: Dissecting Molecular Circuits with Scalable Single-Cell RNA Profiling of Pooled Genetic Screens.” Cell 167 (7): 1853–66.e17.

Domingo, Júlia, Mariia Minaeva, John A. Morris, Marcello Ziosi, Neville E. Sanjana, and Tuuli Lappalainen. 2024. “Non-Linear Transcriptional Responses to Gradual Modulation of Transcription Factor Dosage.” bioRxiv : The Preprint Server for Biology, March. 10.1101/2024.03.01.582837.

Feldman, David, Avtar Singh, Jonathan L. Schmid-Burgk, Rebecca J. Carlson, Anja Mezger, Anthony J. Garrity, Feng Zhang, and Paul C. Blainey. 2019. “Optical Pooled Screens in Human Cells.” Cell 179 (3): 787–99.e17.

Finucane, Hilary K., Yakir A. Reshef, Verneri Anttila, Kamil Slowikowski, Alexander Gusev, Andrea Byrnes, Steven Gazal, et al. 2018. “Heritability Enrichment of Specifically Expressed Genes Identifies Disease-Relevant Tissues and Cell Types.” Nature Genetics 50 (4): 621–29.

Gandal, Michael J., Jillian R. Haney, Neelroop N. Parikshak, Virpi Leppa, Gokul Ramaswami, Chris Hartl, Andrew J. Schork, et al. 2018. “Shared Molecular Neuropathology across Major Psychiatric Disorders Parallels Polygenic Overlap.” Science 359 (6376): 693–97.

Gu, Jiacheng, Abhishek Iyer, Ben Wesley, Angelo Taglialatela, Giuseppe Leuzzi, Sho Hangai, Aubrianna Decker, et al. 2023. “CRISPRmap: Sequencing-Free Optical Pooled Screens Mapping Multi-Omic Phenotypes in Cells and Tissue.” *bioRxiv : The Preprint Server for Biology*, December. 10.1101/2023.12.26.572587.

Jaitin, Diego Adhemar, Assaf Weiner, Ido Yofe, David Lara-Astiaso, Hadas Keren-Shaul, Eyal David, Tomer Meir Salame, Amos Tanay, Alexander van Oudenaarden, and Ido Amit. 2016. “Dissecting Immune Circuits by Linking CRISPR-Pooled Screens with Single-Cell RNA-Seq.” Cell 167 (7): 1883–96.e15.

Ji, Yuge, Tessa D. Green, Stefan Peidli, Mojtaba Bahrami, Meiqi Liu, Luke Zappia, Karin Hrovatin, Chris Sander, and Fabian J. Theis. 2023. “Optimal Distance Metrics for Single-Cell RNA-Seq Populations.” bioRxiv. 10.1101/2023.12.26.572833.

Jost, Marco, Daniel A. Santos, Reuben A. Saunders, Max A. Horlbeck, John S. Hawkins, Sonia M. Scaria, Thomas M. Norman, et al. 2020. “Titrating Gene Expression Using Libraries of Systematically Attenuated CRISPR Guide RNAs.” Nature Biotechnology 38 (3): 355–64.

Kang, J. B., A. Z. Shen, S. Gurajala, A. Nathan, L. Rumker, V. R. C. Aguiar, C. Valencia, et al. 2023. “Mapping the Dynamic Genetic Regulatory Architecture of HLA Genes at Single-Cell Resolution.” Nature Genetics 55 (12). 10.1038/s41588-023-01586-6.

Karczewski, Konrad J., Laurent C. Francioli, Grace Tiao, Beryl B. Cummings, Jessica Alföldi, Qingbo Wang, Ryan L. Collins, et al. 2020. “The Mutational Constraint Spectrum Quantified from Variation in 141,456 Humans.” Nature 581 (7809): 434–43.

Kudo, Takamasa, Ana M. Meireles, Reuben Moncada, Yushu Chen, Ping Wu, Joshua Gould, Xiaoyu Hu, et al. 2023. “Highly Multiplexed, Image-Based Pooled Screens in Primary Cells and Tissues with PerturbView.” bioRxiv. 10.1101/2023.12.26.573143.

Lacher, Sonja M., Julia Bruttger, Bettina Kalt, Jean Berthelet, Krishnaraj Rajalingam, Simone Wörtge, and Ari Waisman. 2017. “HMG-CoA Reductase Promotes Protein Prenylation and Therefore Is Indispensible for T-Cell Survival.” Cell Death & Disease 8 (5): e2824.

Lee, S. H., J. Yang, M. E. Goddard, P. M. Visscher, and N. R. Wray. 2012. “Estimation of Pleiotropy between Complex Diseases Using Single-Nucleotide Polymorphism-Derived Genomic Relationships and Restricted Maximum Likelihood.” Bioinformatics 28 (19): 2540–42.

Lin, Yingxin, Shila Ghazanfar, Dario Strbenac, Andy Wang, Ellis Patrick, David M. Lin, Terence Speed, Jean Y. H. Yang, and Pengyi Yang. 2019. “Evaluating Stably Expressed Genes in Single Cells.” GigaScience 8 (9). 10.1093/gigascience/giz106.

Lopez, Romain, Jeffrey Regier, Michael B. Cole, Michael I. Jordan, and Nir Yosef. 2018. “Deep Generative Modeling for Single-Cell Transcriptomics.” Nature Methods 15 (12): 1053–58.

Love, Michael I., Wolfgang Huber, and Simon Anders. 2014. “Moderated Estimation of Fold Change and Dispersion for RNA-Seq Data with DESeq2.” Genome Biology 15 (12): 1–21.

Ma, Ying, Shiquan Sun, Xuequn Shang, Evan T. Keller, Mengjie Chen, and Xiang Zhou. 2020. “Integrative Differential Expression and Gene Set Enrichment Analysis Using Summary Statistics for scRNA-Seq Studies.” Nature Communications 11 (1): 1585.

Meyers, Robin M., Jordan G. Bryan, James M. McFarland, Barbara A. Weir, Ann E. Sizemore, Han Xu, Neekesh V. Dharia, et al. 2017. “Computational Correction of Copy Number Effect Improves Specificity of CRISPR–Cas9 Essentiality Screens in Cancer Cells.” Nature Genetics 49 (12): 1779–84.

Morris, John A., Jennifer S. Sun, and Neville E. Sanjana. 2024. “Next-Generation Forward Genetic Screens: Uniting High-Throughput Perturbations with Single-Cell Analysis.” Trends in Genetics: TIG 40 (2): 118–33.

Nabet, Behnam, Justin M. Roberts, Dennis L. Buckley, Joshiawa Paulk, Shiva Dastjerdi, Annan Yang, Alan L. Leggett, et al. 2018. “The dTAG System for Immediate and Target-Specific Protein Degradation.” Nature Chemical Biology 14 (5): 431–41.

Naqvi, Sahin, Seungsoo Kim, Hanne Hoskens, Harold S. Matthews, Richard A. Spritz, Ophir D. Klein, Benedikt Hallgrímsson, et al. 2023. “Precise Modulation of Transcription Factor Levels Identifies Features Underlying Dosage Sensitivity.” Nature Genetics 55 (5): 841–51.

O’Connor, Luke J., Armin P. Schoech, Farhad Hormozdiari, Steven Gazal, Nick Patterson, and Alkes L. Price. 2019. “Extreme Polygenicity of Complex Traits Is Explained by Negative Selection.” American Journal of Human Genetics 105 (3): 456–76.

Peidli, Stefan, Tessa D. Green, Ciyue Shen, Torsten Gross, Joseph Min, Samuele Garda, Bo Yuan, et al. 2024. “scPerturb: Harmonized Single-Cell Perturbation Data.” Nature Methods 21 (3): 531–40.

Plaisier, Seema B., Richard Taschereau, Justin A. Wong, and Thomas G. Graeber. 2010. “Rank–rank Hypergeometric Overlap: Identification of Statistically Significant Overlap between Gene-Expression Signatures.” Nucleic Acids Research 38 (17): e169.

Pratapa, Aditya, Amogh P. Jalihal, Jeffrey N. Law, Aditya Bharadwaj, and T. M. Murali. 2020. “Benchmarking Algorithms for Gene Regulatory Network Inference from Single-Cell Transcriptomic Data.” Nature Methods 17 (2): 147–54.

Replogle, Joseph M., Reuben A. Saunders, Angela N. Pogson, Jeffrey A. Hussmann, Alexander Lenail, Alina Guna, Lauren Mascibroda, et al. 2022. “Mapping Information-Rich Genotype-Phenotype Landscapes with Genome-Scale Perturb-Seq.” Cell 185 (14): 2559–75.e28.

Reshef, Yakir A., Laurie Rumker, Joyce B. Kang, Aparna Nathan, Ilya Korsunsky, Samira Asgari, Megan B. Murray, D. Branch Moody, and Soumya Raychaudhuri. 2022. “Co-Varying Neighborhood Analysis Identifies Cell Populations Associated with Phenotypes of Interest from Single-Cell Transcriptomics.” Nature Biotechnology 40 (3): 355–63.

Rood, Jennifer E., Aidan Maartens, Anna Hupalowska, Sarah A. Teichmann, and Aviv Regev. 2022. “Impact of the Human Cell Atlas on Medicine.” Nature Medicine 28 (12): 2486–96.

Rubin, Adam J., Kevin R. Parker, Ansuman T. Satpathy, Yanyan Qi, Beijing Wu, Alvin J. Ong, Maxwell R. Mumbach, et al. 2019. “Coupled Single-Cell CRISPR Screening and Epigenomic Profiling Reveals Causal Gene Regulatory Networks.” Cell 176 (1-2): 361–76.e17.

Simmons, Sean K., Gila Lithwick-Yanai, Xian Adiconis, Florian Oberstrass, Nika Iremadze, Kathryn Geiger-Schuller, Pratiksha I. Thakore, et al. 2023. “Mostly Natural Sequencing-by-Synthesis for scRNA-Seq Using Ultima Sequencing.” Nature Biotechnology 41 (2): 204–11.

Stephens, Matthew. 2016. “False Discovery Rates: A New Deal.” Biostatistics 18 (2): 275–94.

Subramanian, Aravind, Pablo Tamayo, Vamsi K. Mootha, Sayan Mukherjee, Benjamin L. Ebert, Michael A. Gillette, Amanda Paulovich, et al. 2005. “Gene Set Enrichment Analysis: A Knowledge-Based Approach for Interpreting Genome-Wide Expression Profiles.” Proceedings of the National Academy of Sciences of the United States of America 102 (43): 15545–50.

Weber, Christopher M., Antonina Hafner, Jacob G. Kirkland, Simon M. G. Braun, Benjamin Z. Stanton, Alistair N. Boettiger, and Gerald R. Crabtree. 2021. “mSWI/SNF Promotes Polycomb Repression Both Directly and through Genome-Wide Redistribution.” Nature Structural & Molecular Biology 28 (6): 501–11.

Weiss, M. J., G. Keller, and S. H. Orkin. 1994. “Novel Insights into Erythroid Development Revealed through in Vitro Differentiation of GATA-1 Embryonic Stem Cells.” Genes & Development 8 (10): 1184–97.

Xu, Zihan, Andras Sziraki, Jasper Lee, Wei Zhou, and Junyue Cao. 2023. “Dissecting Key Regulators of Transcriptome Kinetics through Scalable Single-Cell RNA Profiling of Pooled CRISPR Screens.” *Nature Biotechnology*, September. 10.1038/s41587-023-01948-9.

Yang, Jian, Beben Benyamin, Brian P. McEvoy, Scott Gordon, Anjali K. Henders, Dale R. Nyholt, Pamela A. Madden, et al. 2010. “Common SNPs Explain a Large Proportion of the Heritability for Human Height.” Nature Genetics 42 (7): 565–69.

Yao, Douglas, Loic Binan, Jon Bezney, Brooke Simonton, Jahanara Freedman, Chris J. Frangieh, Kushal Dey, et al. 2023. “Scalable Genetic Screening for Regulatory Circuits Using Compressed Perturb-Seq.” *Nature Biotechnology*, October. 10.1038/s41587-023-01964-9.

Yazar, Seyhan, Jose Alquicira-Hernandez, Kristof Wing, Anne Senabouth, M. Grace Gordon, Stacey Andersen, Qinyi Lu, et al. 2022. “Single-Cell eQTL Mapping Identifies Cell Type– specific Genetic Control of Autoimmune Disease.” *Science*, April. 10.1126/science.abf3041.

