## Supplementary Figures for "Transcriptome-wide characterization of genetic perturbations"

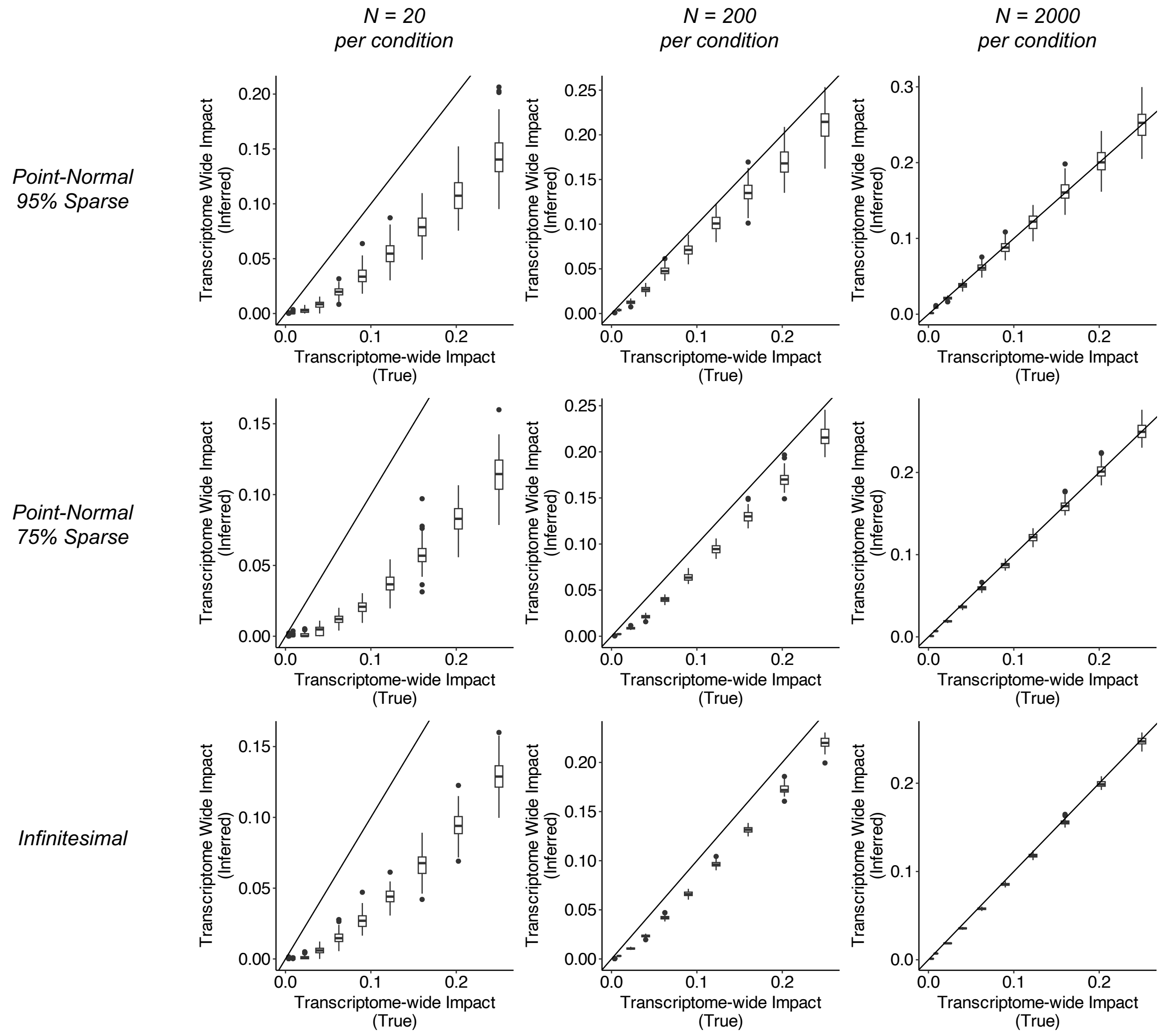


**Supplementary Figure 1: Simulations (Transcriptome-wide impact).** In these simulations, we simulated cell-wise counts using mean and overdispersion parameters inferred from 10 batches of the K562-Essential dataset, and then simulated perturbed counts using various effect size distribution shapes (“Point-Normal 95% Sparse”, “Point Normal 75% Sparse”, “Infinitesimal”, see Methods), sample sizes (20, 200, or 2000 per condition), and transcriptome-wide impacts (i.e. effect size variances). For each simulation, the true transcriptome-wide impact is on the x-axis, and the inferred transcriptome-wide impact is on the y-axis

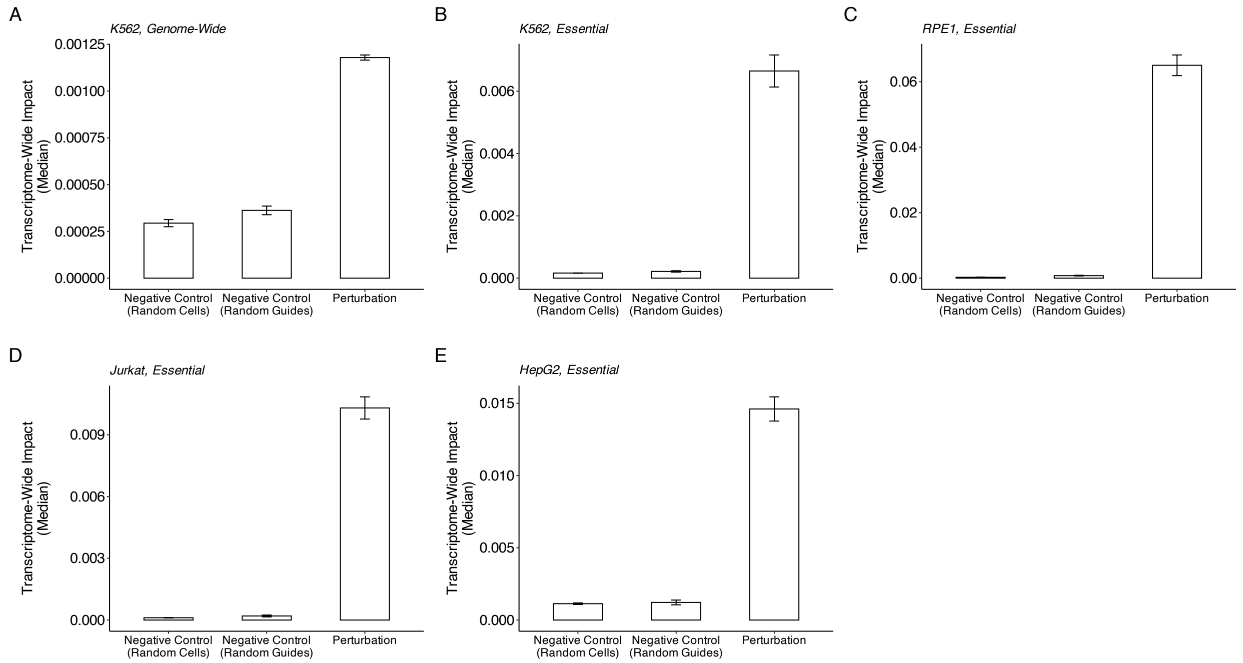


**Supplementary Figure 2: Empirical negative control analyses.** For each experiment, we performed two negative control analyses. In the “Random Cells” analysis, we randomly selected cells carrying non-targeting guide RNAs as “perturbed” cells, and conducted pseudobulk differential expression and TRADE analysis, i.e. including cells from many different non-targeting guides. We did so 500 times for each cell type, drawing the number of “perturbed” cells from the empirical distribution of targeting guide counts for each experiment. For the “Random Guides” analysis, we conducted pseudobulk differential expression and TRADE analysis, for each non-targeting guide, i.e. treating each non-targeting guide as a “perturbation”. For each panel, the median of negative-control TI estimates is shown in comparison with the median of TI estimates for targeting guide RNAs.

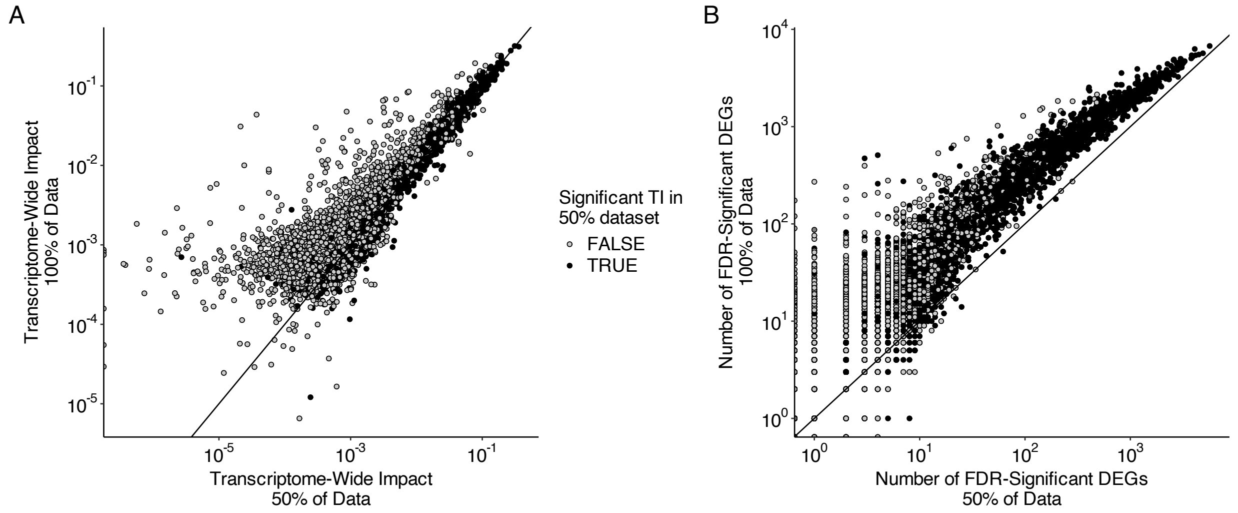


**Supplementary Figure 3: Effects of Downsampling on Transcriptome-wide Impact**
(**A**) To understand why transcriptome-wide impact estimates were slightly smaller in 50% downsampled data, we plotted total cumulative differential expression estimates between the full and downsampled. This revealed that the points far from the y=x line were the points without significant transcriptome-wide impact in the downsampled datasets, where the estimates in the downsampled dataset were greater. This analysis suggests that the subtle decrease in transcriptome-wide impact with downsampling arises from an inability to resolve small nonzero effects after crossing a power threshold. (**B**) In contrast, the number of FDR-significant genes is consistently much smaller after downsampling by 50%.

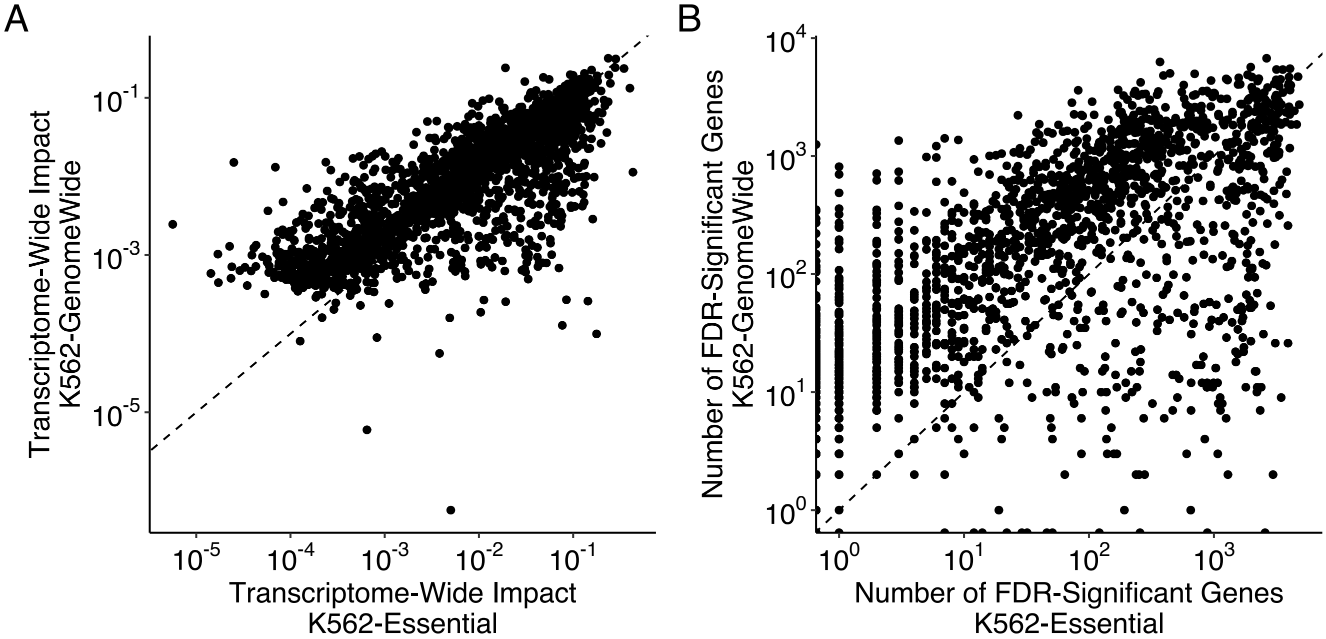


**Supplementary Figure 4: Consistency of Transcriptome-Wide Impact and Number of DEGs.** (A) Comparison of transcriptome-wide impact estimates for perturbation of the same 2,057 essential genes in the K562-Essential and K562-GenomeWide experiments ($R^2$ = 59.7%). (B) Comparison of the number of FDR-significant genes for the same set of perturbations between the K562-Essential and K562-GenomeWide experiments ($R^2$ = 28.4%).

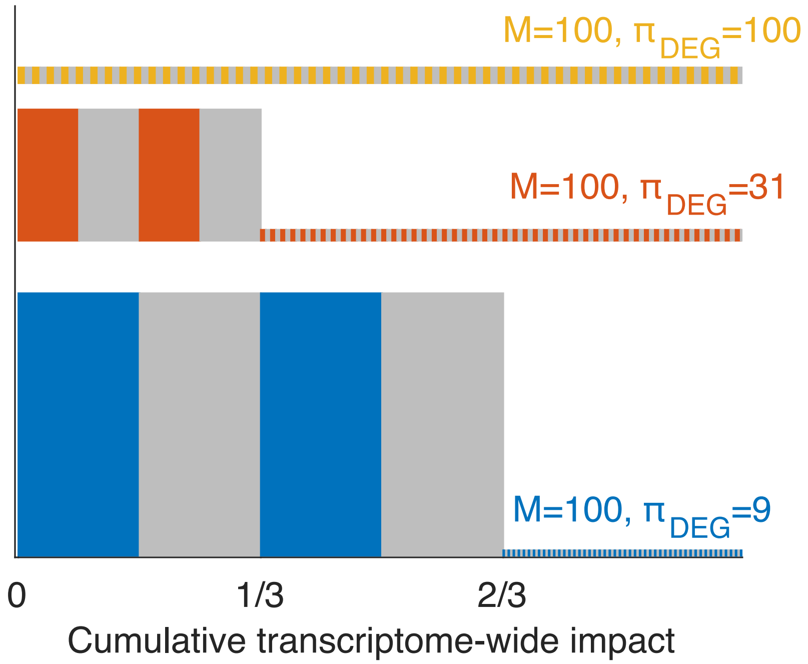


**Supplementary Figure 5:$\pi_{DEG}$, the effective number of differentially expressed genes**. Example distributions of effect sizes and the resulting number of non-null differentially expressed genes (M) and effective number of differentially expressed genes ($\pi_{DEG}$), in a scenario with 100 measured genes. The x-axis shows the cumulative transcriptome-wide impact. In the scenario where all genes have the same effect size (top line, orange), M and $\pi_{DEG}$ are both equal to 100. When a few genes explain a large fraction (middle line, red) or dominate (bottom line, blue) transcriptome-wide impact, but the other genes have small, non-zero effects, $\pi_{DEG}$ captures this qualitatively different architecture, while M is still equal to 100.

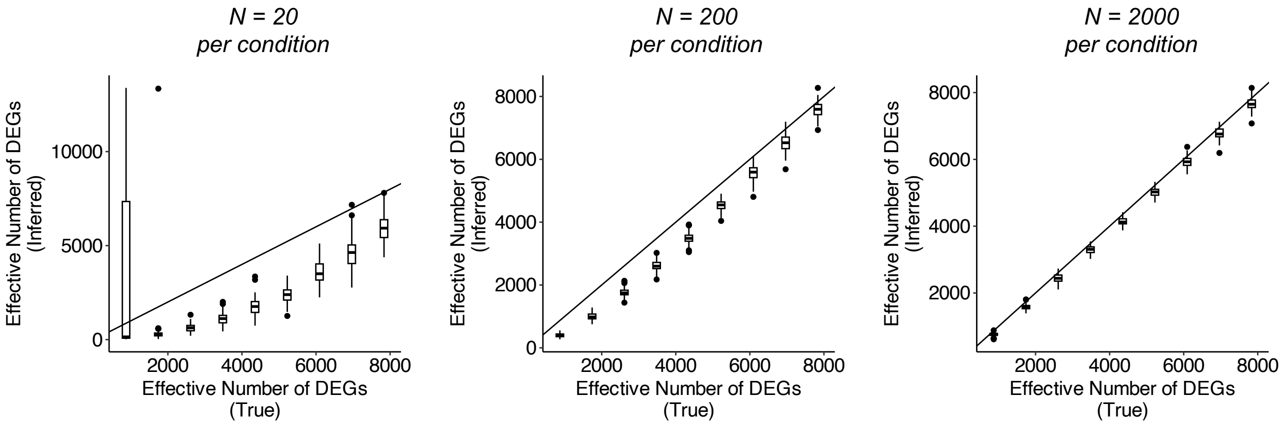


**Supplementary Figure 6: Simulations ($\pi_{DEG}$).** Simulations with a similar structure to those described in Supplementary Figure 1, but with a point normal distribution with varying proportion of non-null effects; this proportion times the number of genes is the effective number of DEGs ($\pi_{DEG}$). As in Supplementary Figure 1, we simulated experiments with N = 20, 200, and 2000 cells per perturbation. On the x axis is the true $\pi_{DEG}$, and the y axis is the $\pi_{DEG}$ inferred with TRADE.

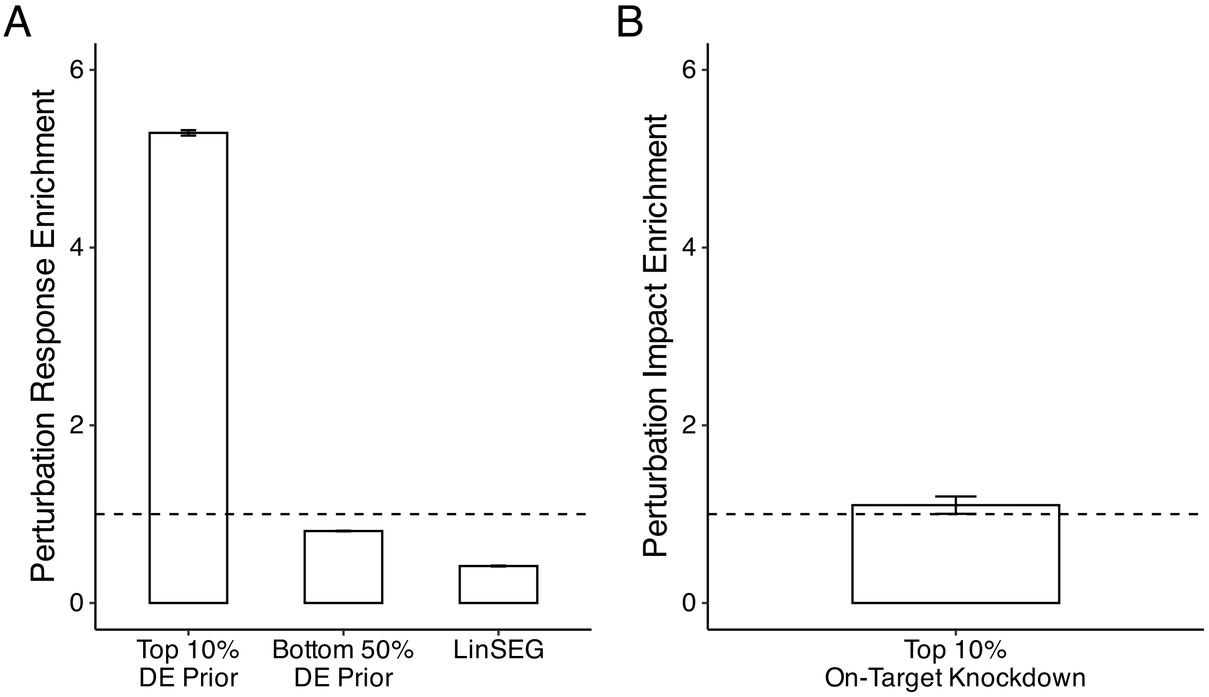


**Supplementary Figure 7: Positive Control Enrichments. (A)** Perturbation response enrichment estimates for three gene-sets expected to be enriched or depleted for differential expression signal, averaged across perturbations in the K562-GenomeWide experiment. “Top 10% DE Prior” and “Bottom 50% DE Prior” are derived from estimates of the DE Prior from Crow et al, 2019, who predicted which genes are more or less likely to be represented in significant differentially expressed gene lists from a database of functional genomic data. “LinSeg” is the set of stably-expressed genes identified by Lin et al (2019), genes which are similar in expression across evolution and development. Point estimate for Top 10% DE Prior: 5.85 (sem = 0.03). Point estimate for Top 50% DE Prior: 0.79 (sem = 0.003). Point estimate for LinSEG: 0.38 (sem = 0.003). **(B)** Perturbation impact enrichment estimate for genes with in the top decile for on-target knockdown, as quantified by the DESeq2 log2FoldChange. Point estimate: 1.1 (sem = 0.1)

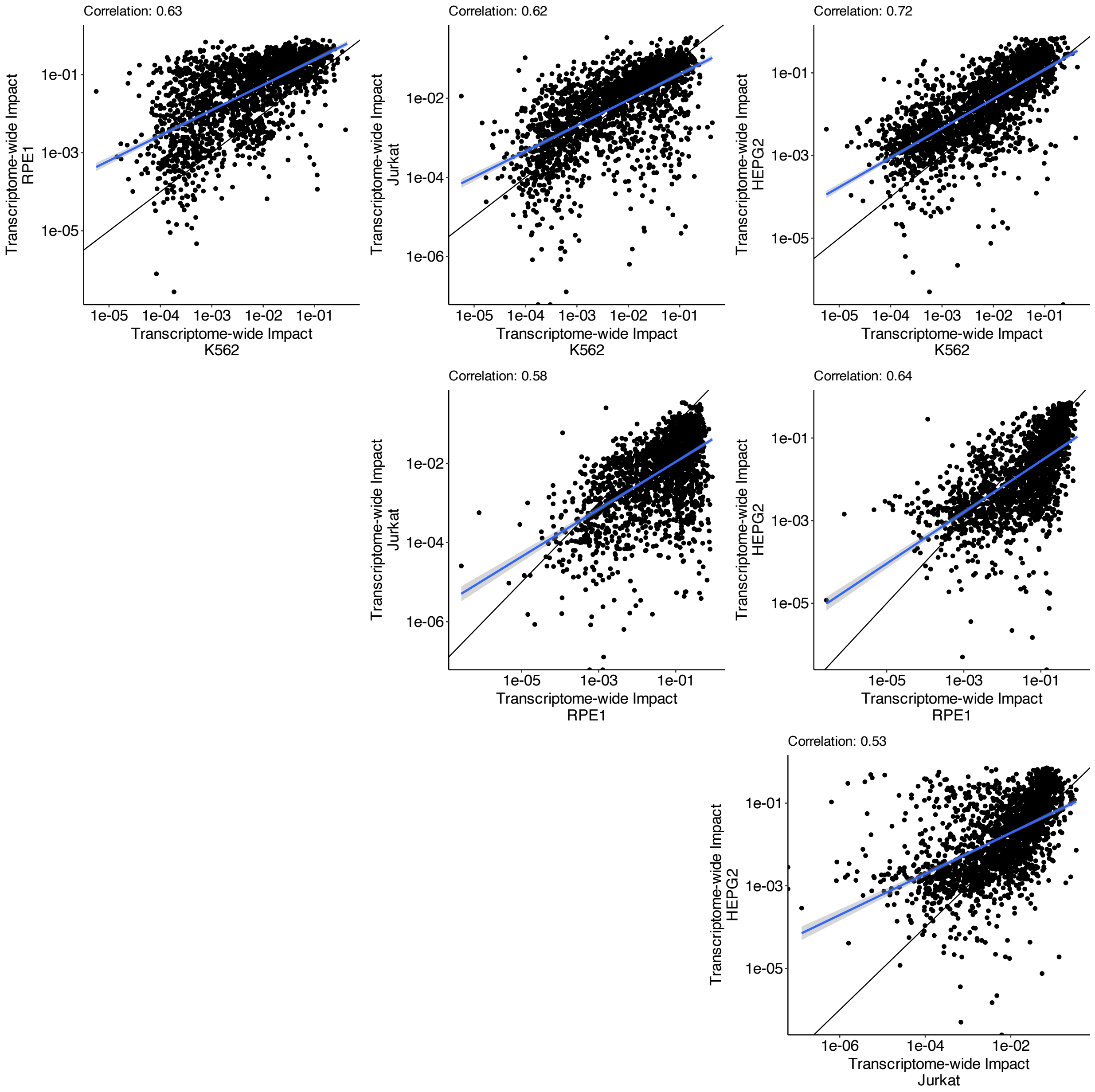


**Supplementary Figure 8: TI Comparison Across Cell Types.** For each pair of cell types, we compared transcriptome-wide impact estimates for common essential perturbations.

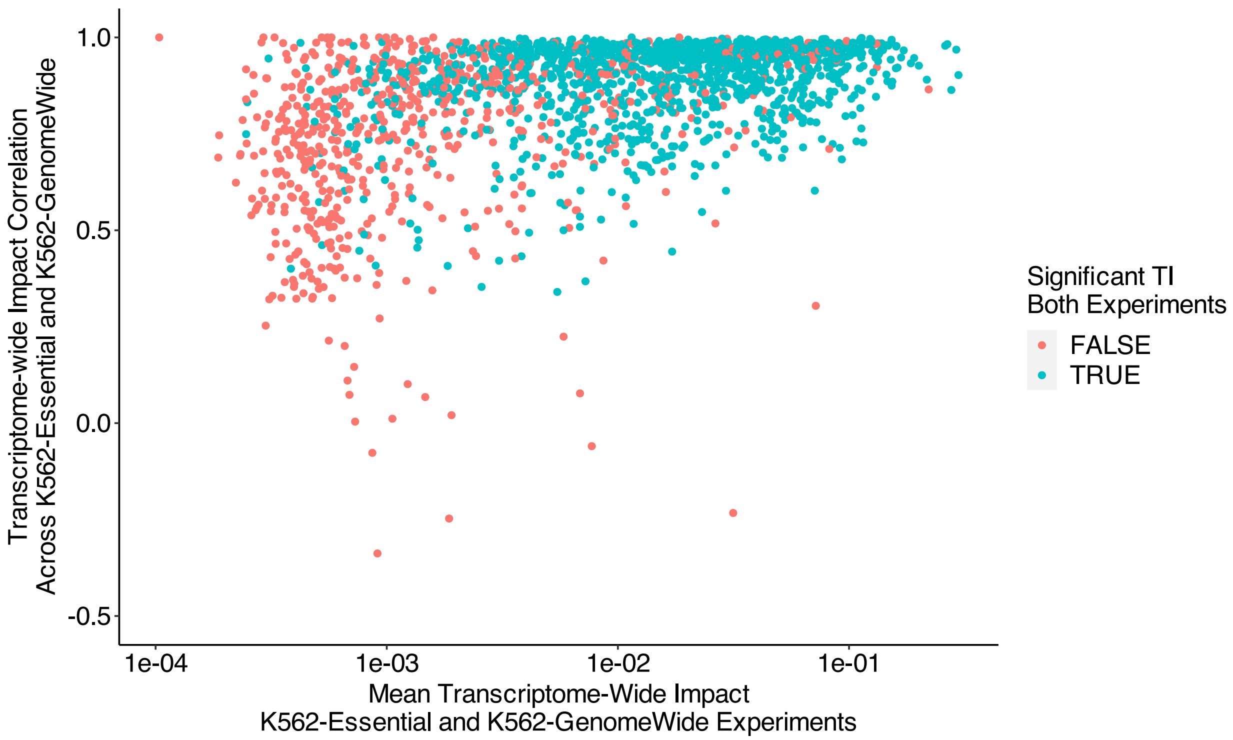


**Supplementary Figure 9: Perturbations with low TRADE replication correlations.** Comparison of transcriptome-wide impact correlation in the K562-Essential and K562-GenomeWide experiments, with mean transcriptome-wide impact in two experiments (i.e., correlation of transcriptomic effects across replicate experiments vs mean of transcriptomic impact across replicate experiments). Color reflects whether transcriptome-wide impact estimates are significant in both experiments.

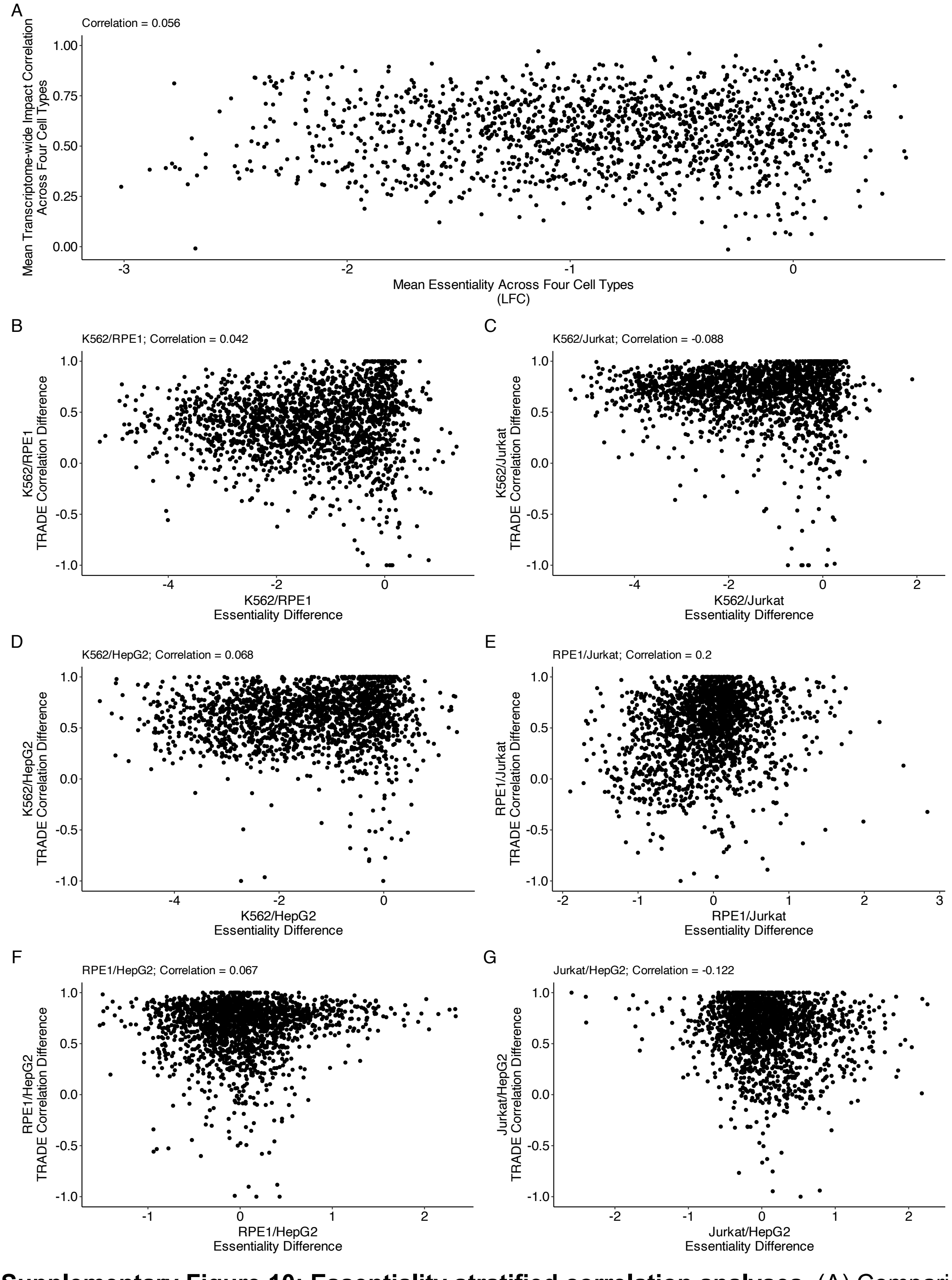


**Supplementary Figure 10: Essentiality stratified correlation analyses**. (A) Comparison of mean average growth effect (i.e. quantitative essentiality) of common essential gene perturbations, and correlation of transcriptomic effects averaged across all six cell-type pairs. (B - G) Comparison of inter-cell-type differences in growth effect/essentiality, with transcriptome-wide impact correlation of perturbation effects, across common essential gene perturbations.

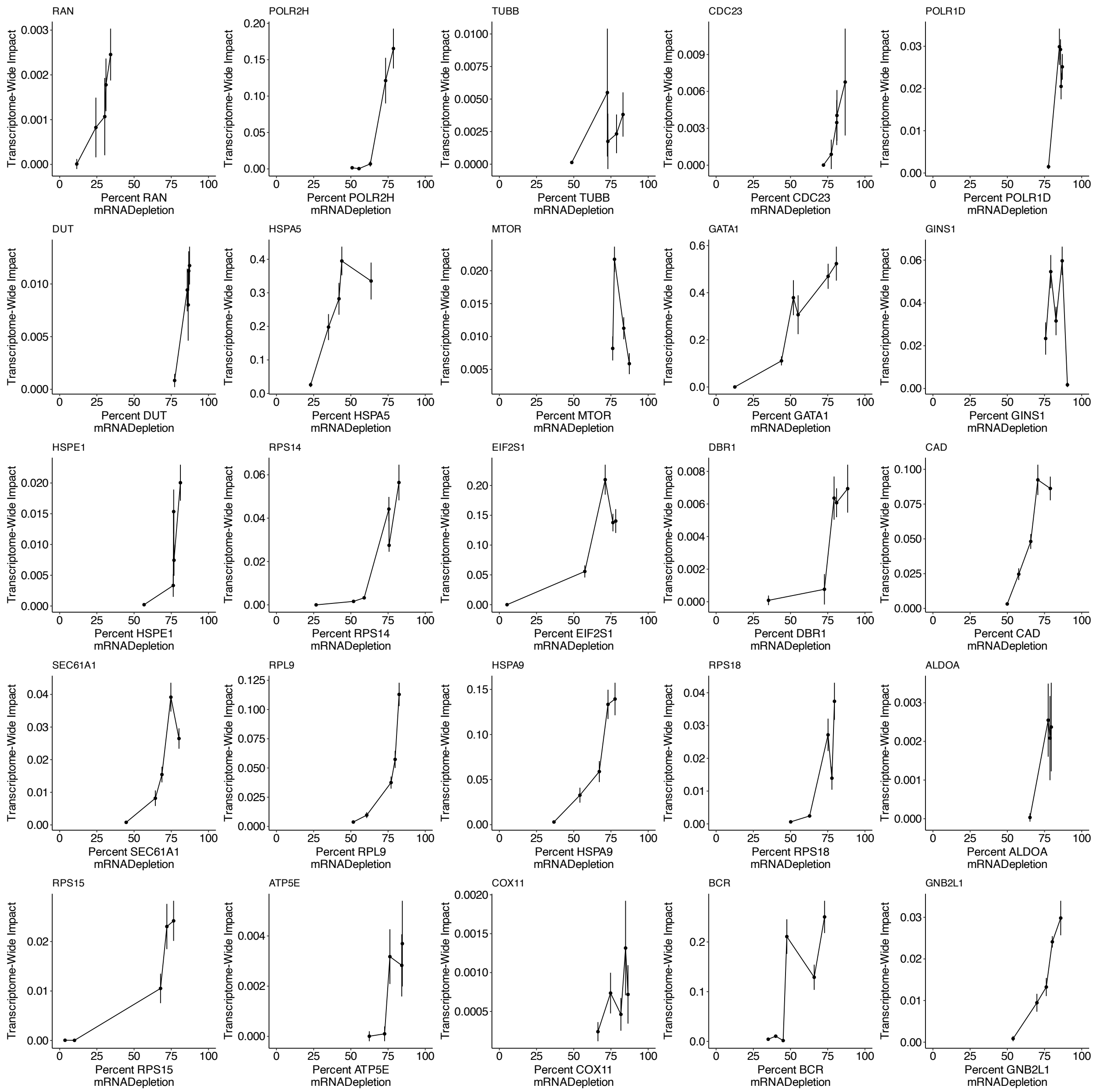


**Supplementary Figure 11: TI curves for Jost et al CRISPR experiment**. For each of the 25 essential gene perturbations in K562 profiled by Jost et al (2020), comparison of degree of target knockdown (estimated from single-cell RNA-seq) with transcriptome-wide impact of perturbation. Note that y axis scales differ between plots.

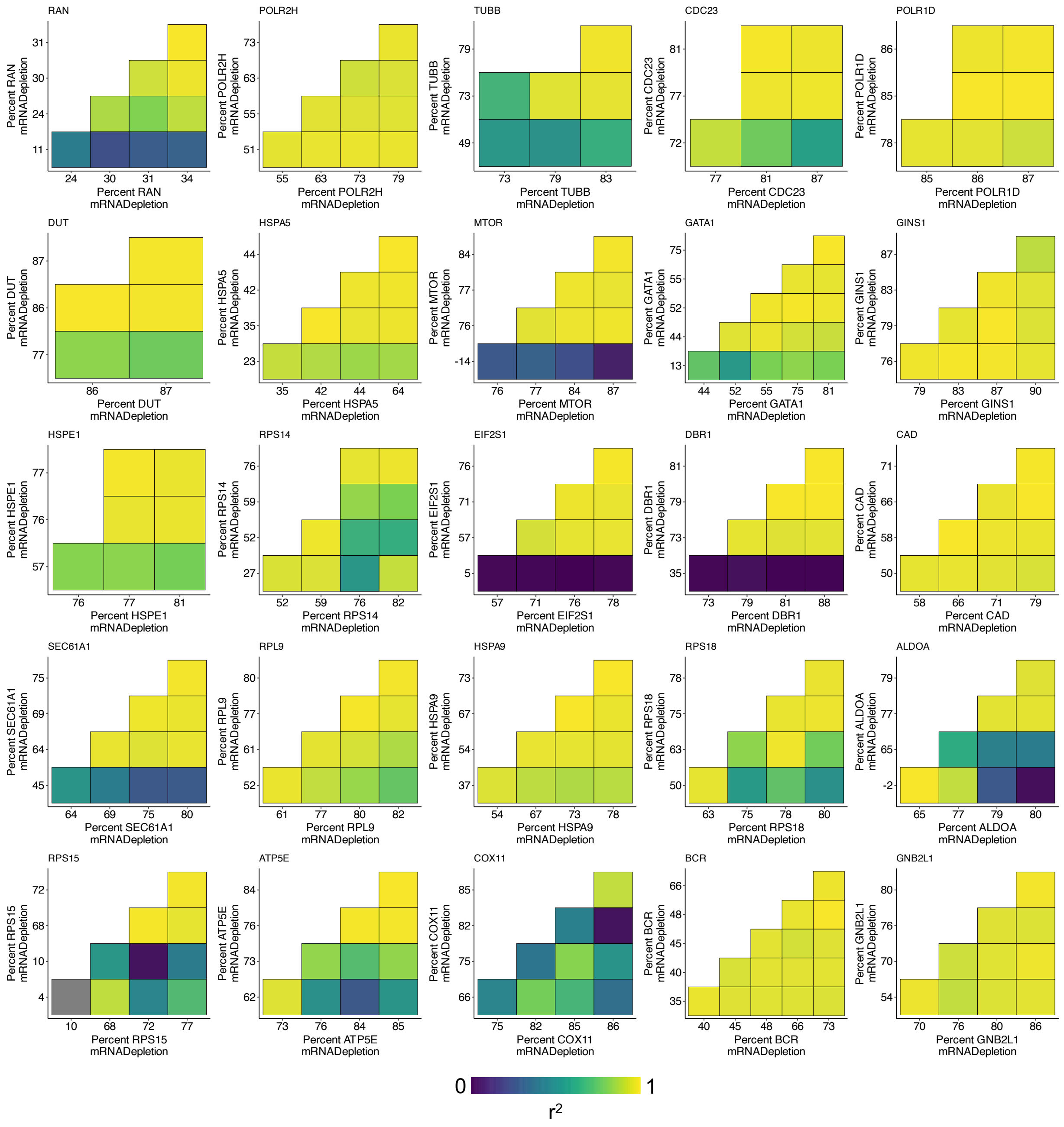


**Supplementary Figure 12: TRADE correlation matrices for Jost et al CRISPR experiment.** For each of the 25 essential gene perturbations in K562 profiled by Jost et al (2020), estimates of transcriptome-wide impact correlation across different perturbation dosages. Note that both x and y axis labels vary across plots.

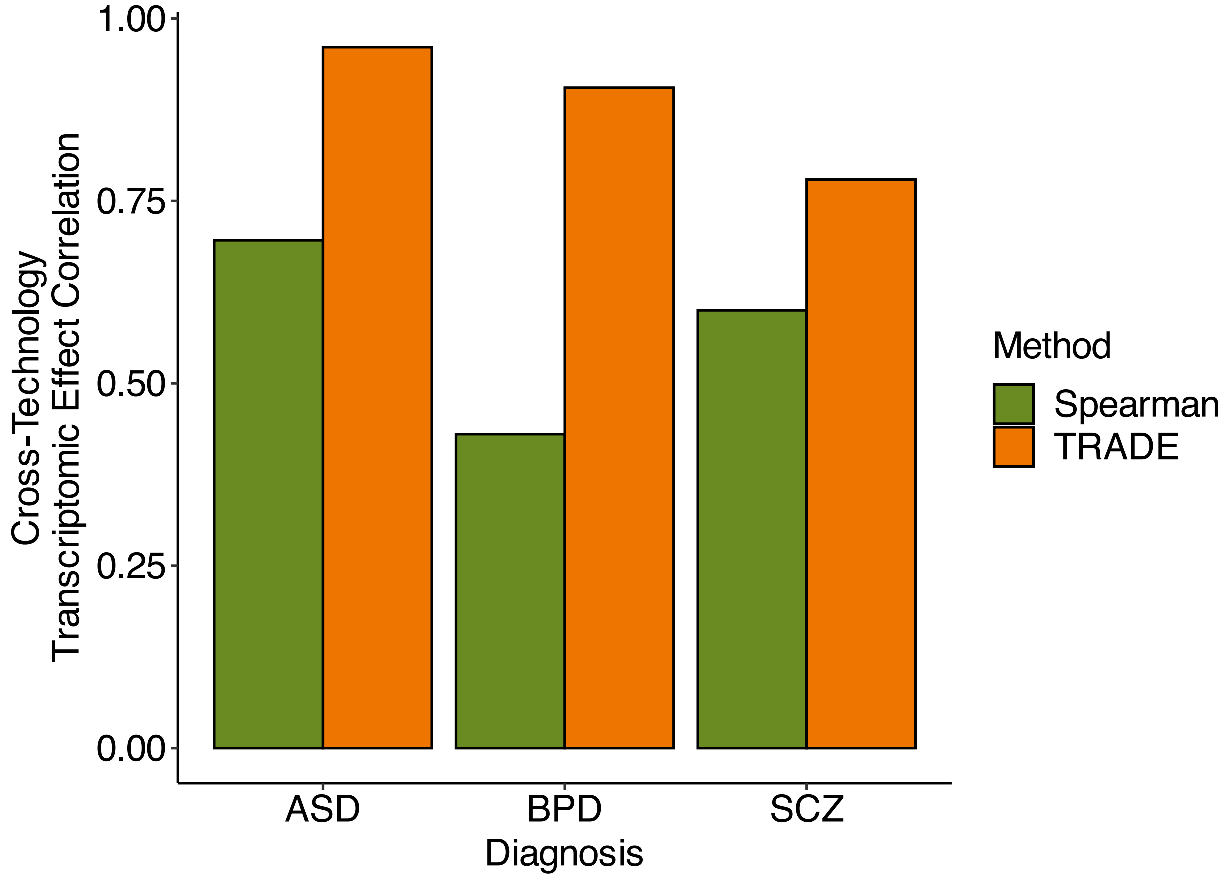


**Supplementary Figure 13: Inter-technology correlations for neuropsychiatric conditions**. Correlation of log2FoldChange estimates between microarray and RNA-Seq datasets from the PsychENCODE resource, estimated with the sample Spearman correlation (green) or TRADE (orange).

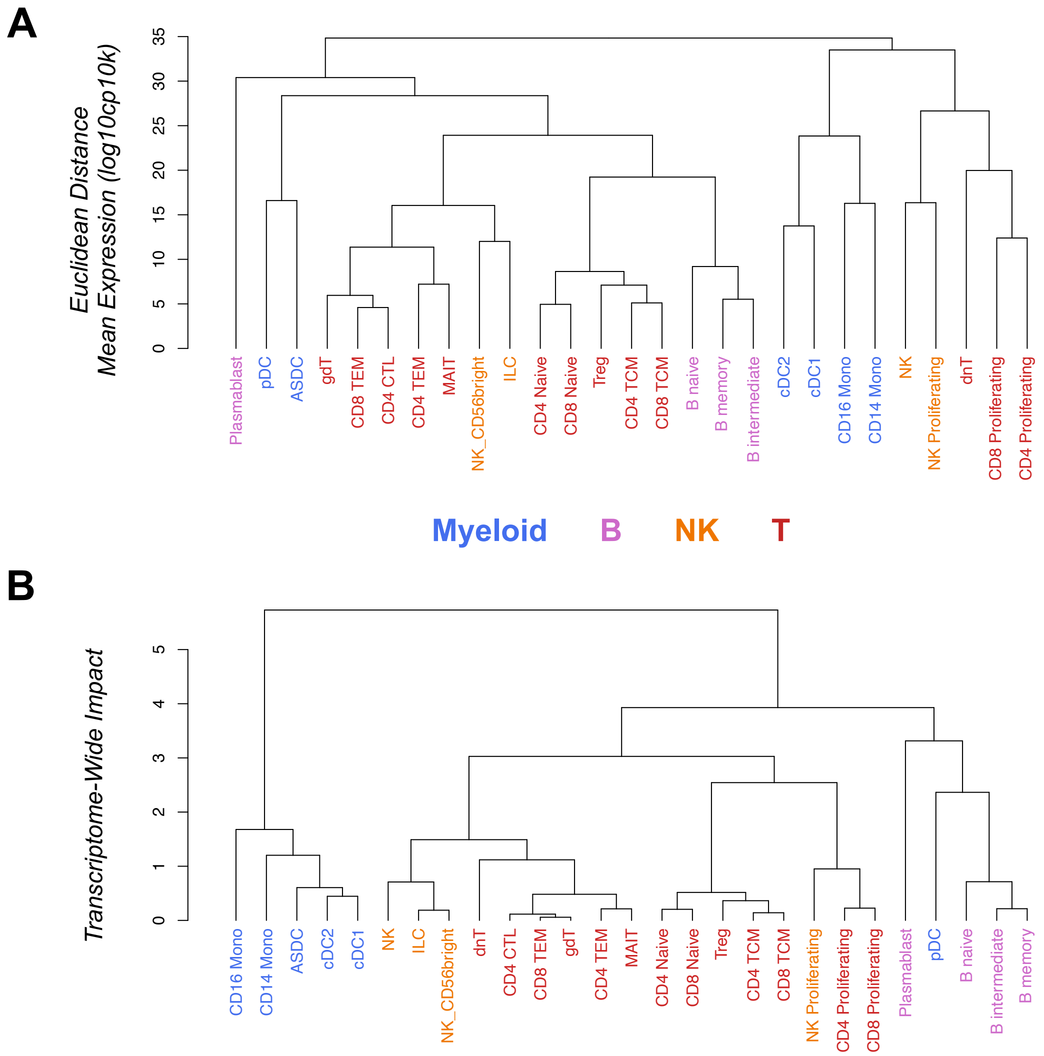


**Supplementary Figure 14 Hierarchically organizing cell-types with TRADE.** We applied TRADE to the OneK1K dataset, which profiles 822,522 PBMCs from 969 donors (**Methods**). We chose this dataset because PBMCs are relatively well annotated, with known functional and lineage relationships between readily identifiable cell types. Compared to Euclidean distance, hierarchically clustering cell types by transcriptome-wide impact as a distance metric lead to an inferred hierarchy that was more biologically plausible. In particular, several lowly sampled cell types were placed in more plausible positions within the cell type hierarchy. For example, plasmablasts (a low-abundance B-cell subtype) were clustered far away from other B-cells via Euclidean distance, but placed much closer to other B-cells via transcriptome-wide impact. Similarly, ILCs and NK_CDK65Bright, two rare subtypes of NK-cells, were placed close to NK cells via transcriptome-wide impact but not Euclidean distance. Moreover, TRADE recapitulated the early developmental split between myeloid and lymphoid cells, whereas Euclidean distance frequently placed both types of cells within the same clade. The only exception to this myeloid/lymphoid divide in the TRADE hierarchy are plasmacytoid dendritic cells (pDCs), which were placed near B-cells; notably, pDCs are named for their morphological similarity to B-cells (plasma cells), suggesting that they may truly transcriptomically resemble lymphoid cells.
