## Supplementary Appendices for "Transcriptome-wide characterization of genetic perturbations"

### Supplementary Appendix 1: Transcriptome-wide impact and $\pi_{DEG}$

Transcriptome-wide impact and $\pi_{DEG}$ capture different aspects of the distribution of differential effects. Transcriptome-wide impact is the variance of the effect-size distribution, and $\pi_{DEG}$ is a function of the kurtosis of the effect size distribution, that captures the "effective number of differentially expressed genes". The relationship between transcriptome-wide impact and $\pi_{DEG}$ gives insight into how effects are spread across genes.

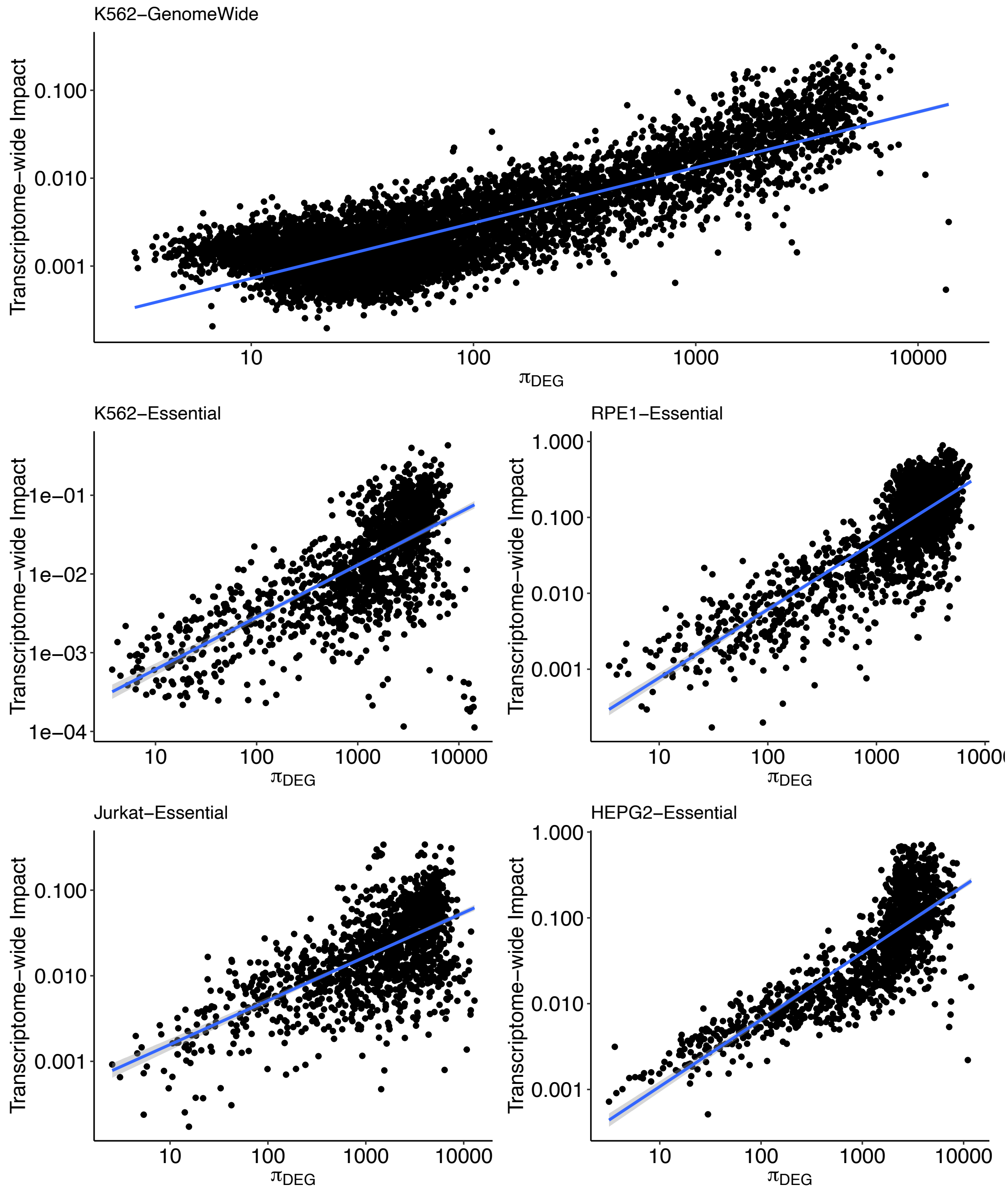


(Notably, this and the below analyses are restricted to perturbations that caused significant transcriptome-wide impact, due to the instability of $\pi_{DEG}$ in the setting of low/non-significant transcriptome-wide impact)

As predicted, transcriptome-wide impact and $\pi_{DEG}$ are positively associated; as the number of genes affected rises, so too does the amount of differential expression signal. Less trivially, there appear to be no perturbations that achieve a large transcriptome-wide impact while affecting a few genes (i.e. paucity of points in the top-left of the above plots). This suggests that no perturbations cause large transcriptomic change via very large effects on a small set of genes; rather, when perturbations cause large transcriptomic change, they do so by affecting many genes.

A typical log2(FoldChange) magnitude of affected genes $\sigma$ can be computed with the ratio of transcriptome-wide impact to $\pi_{DEG}$:

$$\sigma = \sqrt{\frac{n_{genes} * TI}{\pi_{DEG}}}$$

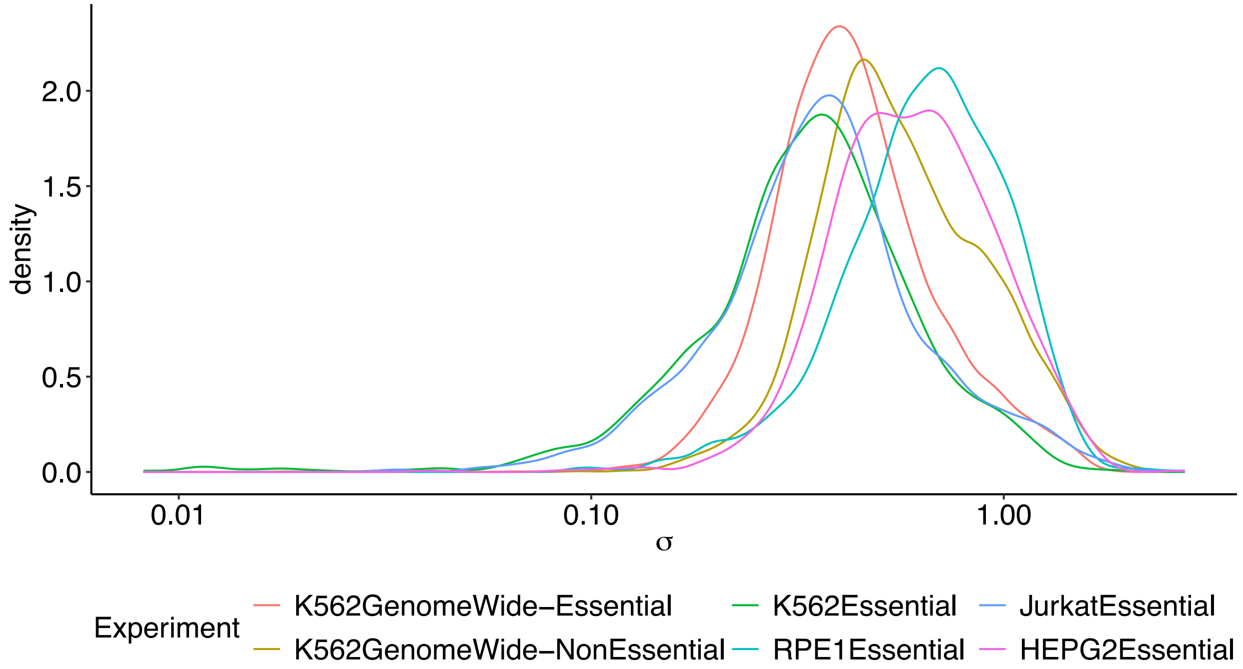


Examining the distribution of $\sigma$ reveals several aspects of perturbation effect architecture:

- For all experiments, the variability in $\sigma$ is mostly limited to the range (0.1,1), suggesting that typical differential expression effects exhibit a limited range across perturbations
- For the K562 cell line, $\sigma$ is larger in the GenomeWide experiment than in the Essential experiment, suggesting that while essential perturbations affect more genes, they do so with a smaller typical effect size
- The mean $\sigma$ varies between cell types. In particular, it is largest in the RPE1 experiments, slightly smaller in the HepG2 experiments, and smaller in the K562 and Jurkat experiments

**Supplementary Appendix 2: Bias of the sample correlation coefficient**

This phenomenon, known as *attenuation*, was identified in Spearman (1904); the arguments from that paper are paraphrased here.

Consider two sets of effect sizes (i.e. log2FoldChanges) that are estimated with additive, independent measurement error:

$$\widehat{\beta_1} = \beta_1 + \epsilon_1$$
$$\widehat{\beta_2} = \beta_2 + \epsilon_2$$

The correlation between the estimated effect sizes is:

$$Corr(\widehat{\beta_1}, \widehat{\beta_2}) = \frac{Cov(\widehat{\beta_1}, \widehat{\beta_2})}{\sqrt{Var(\widehat{\beta_1})Var(\widehat{\beta_2})}}$$

$$= \frac{Cov(\beta_1 + \epsilon_1, \beta_2 + \epsilon_2)}{\sqrt{Var(\beta_1 + \epsilon_1)Var(\beta_2 + \epsilon_2)}}$$

$$= \frac{Cov(\beta_1, \beta_2) + Cov(\beta_1, \epsilon_2) + Cov(\epsilon_1, \beta_2) + Cov(\epsilon_1, \epsilon_2)}{\sqrt{Var(\beta_1 + \epsilon_1)Var(\beta_2 + \epsilon_2)}}$$

$$= \frac{Cov(\beta_1, \beta_2)}{\sqrt{Var(\beta_1 + \epsilon_1)Var(\beta_2 + \epsilon_2)}} < \frac{Cov(\beta_1, \beta_2)}{\sqrt{Var(\beta_1)Var(\beta_2)}} = Corr(\beta_1, \beta_2)$$

Therefore, the correlation of the estimated effect sizes is a downwardly biased estimator of true effect size correlation, and this downward bias is related to the relative magnitude of the sampling noise. If the sampling variation is of similar or greater magnitude than the true effect size variance, this bias will be severe.

**Supplementary Appendix 3: Consistency of dose-response curves**

Consider the relationship between the expression of target genes of a gene perturbation. For each target gene $g$, the expression of the target gene at dosage 1 of the perturbed gene is equal to the expression of the target gene at full dosage of the perturbed gene times a function $f_g(dosage)$, that describes the response kinetics of $g$ in response to changing dosage of the perturbed gene.

$$\mathrm{Expression_g(dosage_1)} = f_g(\mathrm{dosage_1}) * \mathrm{Expression_g(full\ dosage)}$$
$$\mathrm{Expression_g(dosage_2)} = f_g(\mathrm{dosage_2}) * \mathrm{Expression_g(full\ dosage)}$$

Rearranging:

$$\mathrm{FoldChange_g(dosage_1)} = f_g(\mathrm{dosage_1})$$
$$\mathrm{FoldChange_g(dosage_2)} = f_g(\mathrm{dosage_2})$$

$$\mathrm{FoldChange_g(dosage_1)} = \frac{f_g(\mathrm{dosage_1})}{f_g(\mathrm{dosage_2})}\mathrm{FoldChange_g(dosage_2)}$$

When $f_g$ is identical across target genes up to a scaling factor, then $\frac{f_g(\mathrm{dosage_1})}{f_g(\mathrm{dosage_2})}$ is constant, and $\mathrm{FoldChange_g(dosage_1)}$ and $\mathrm{FoldChange_g(dosage_2)}$ are perfectly correlated across genes. However, when $f_g$ varies across genes, and $\frac{f_g(\mathrm{dosage_1})}{f_g(\mathrm{dosage_2})}$ is no longer constant, there is an imperfect correlation between $\mathrm{FoldChange_g(dosage_1)}$ and $\mathrm{FoldChange_g(dosage_2)}$. This reveals that consistency of perturbation effects across dosages is at least partially a function of the consistency of downstream response kinetics across target genes.

**Supplementary Appendix 4: Bias of the sample Euclidean Distance**

This phenomenon is described in Mahalanobis (1936), and paraphrased here. It is closely related to the bias of the sample correlation in the presence of uncorrelated noise.

Consider the Euclidean distance between expression states of two clusters of cells:

$$\text{Euclidean Distance}^2 = \sum_g (\mu_g - \nu_g)^2$$

Where $\mu_g$ is the mean expression of gene $g$ in cluster 1, and $\nu_g$ is the mean expression of gene $g$ in cluster 2.

In computing distance metrics for analyses such as hierarchical clustering, this quantity is typically estimated using sample means, which differ from the population mean due to sampling variation with a known distribution:

$$\widehat{\mu_g} = \mu_g + \epsilon_g \qquad \epsilon_g \sim N(0, \frac{\sigma_g^2}{n_1})$$

$$\widehat{\nu_g} = \nu_g + \delta_g \qquad \delta_g \sim N(0, \frac{\tau_g^2}{n_2})$$

Computing the Euclidean Distance with sample means rather than population means, we have:

$$\text{Euclidean Distance}^2 = \sum_g \left(\mu_g + \epsilon_g - (\nu_g + \delta_g)\right)^2$$

Taking expectation, and assuming that $\epsilon_g$ and $\delta_g$ are independent:

$$\sum_g (\mu_g - \nu_g)^2 + \sum_g Var(\epsilon_g) + Var(\delta_g)$$
$$= \sum_g (\mu_g - \nu_g)^2 + \sum_g \frac{\sigma_g^2}{n_1} + \frac{\tau_g^2}{n_2}$$
$$= \text{Euclidean Distance}^2 + \frac{1}{n_1}\sum_g \sigma_g^2 + \frac{1}{n_2}\sum_g \tau_g^2$$

We see that that when estimating Euclidean Distance with sample means, the resulting estimates are upwardly biased in the following manner:

- The upward bias becomes smaller as the number of cells in each cluster ($n_1$ and $n_2$) increases
- The upward bias becomes larger as the variance in expression between cells ($\sigma_g^2$ and $\tau_g^2$) grows.
- The upward bias becomes larger as more genes are included in the calculation.
